# Absence of chromosome axis proteins recruitment prevents meiotic recombination chromosome-wide in the budding yeast *Lachancea kluyveri*

**DOI:** 10.1101/2023.08.28.555118

**Authors:** Sylvain Legrand, Asma Saifudeen, Hélène Bordelet, Julien Vernerey, Arnaud Guille, Amaury Bignaut, Agnès Thierry, Laurent Acquaviva, Maxime Gaudin, Aurore Sanchez, Dominic Johnson, Anne Friedrich, Joseph Schacherer, Matthew Neale, Valérie Borde, Romain Koszul, Bertrand Llorente

**Affiliations:** CRCM, UMR 7258 CNRS, INSERM, Aix Marseille Université, Institut Paoli Calmettes, Marseille, France; Department Genomes and Genetics, Groupe Régulation Spatiale des Génomes, UMR 3525 CNRS, Institut Pasteur, Paris, France; UMR3244 CNRS, Institut Curie, PSL (Paris Sciences and Letters) Research University, Université Pierre et Marie Curie (UPMC), Paris, France; Genome Damage and Stability Centre, University of Sussex, UK; Université de Strasbourg, CNRS, GMGM UMR 7156, Strasbourg, France

## Abstract

Meiotic recombination shows broad variations across species and along chromosomes, and is often suppressed at and around genomic regions determining sexual compatibility such as mating type loci in fungi. Here we show that the absence of Spo11-DSBs and meiotic recombination on Lakl0C-left, the chromosome arm containing the sex locus of the *Lachancea kluyveri* budding yeast, results from the absence of recruitment of the two chromosome axis proteins Red1 and Hop1, essential for proper Spo11-DSBs formation. Furthermore, cytological observation of spread pachytene meiotic chromosomes reveals that Lakl0C-left does not undergo synapsis. However, we show that the behavior of Lakl0C-left is independent of its particularly early replication timing and is not accompanied by any peculiar chromosome structure as detectable by Hi-C in this yet poorly studied yeast. Finally, we observed an accumulation of heterozygous mutations on Lakl0C-left and a sexual dimorphism of the haploid meiotic offspring, supporting a direct effect of this absence of meiotic recombination on *L. kluyveri* genome evolution and fitness. Because suppression of meiotic recombination on sex chromosomes is widely observed across eukaryotes, the novel mechanism for recombination suppression described here may apply to other species, with the potential to impact sex chromosome evolution.

## Introduction

DNA double strand breaks (DSBs) initiate recombination between homologous chromosomes during meiotic prophase, an important event that promotes chromosome segregation, and hence fertility and genome evolution (1–3). The type II topoisomerase-related protein Spo11 is the catalytic subunit of a multiprotein machinery that generates the meiotic DSBs (4–8). Such Spo11-DSBs occur within nucleosome-free DNA, typically found in gene promoters that constitute Spo11-DSB hotspots in the yeast model *Saccharomyces cerevisiae* (9–11). The Spo11-DSB hotspot landscape is relatively well conserved among *Saccharomyces* species, likely because gene promoters are conserved functional elements that are under selective constraints (12).

In *S. cerevisiae,* meiotic prophase chromosomes undergo compaction to form chromatin loops of ca. 20-100 kb (13–18). These chromatin loops are anchored on a protein axis composed of a coalescence of cohesins, the axis-core protein Red1 and the HORMA domain-containing protein Hop1 (19–22). The current model of axis proteins recruitment is that the meiotic specific kleisin cohesin subunit Rec8 ensures most of Red1 recruitment through direct interaction, and that Red1 then recruits its interacting partner, Hop1 (16, 23). A second parallel pathway relies on a direct Hop1-chromatin interaction within broad genomic islands with high gene density, leading to further recruitment of Red1 (24). Spo11-DSB hotspots are specifically located within chromatin loops, away from the chromosome axis attachment sites at the base of these loops (16). While Rec8 is dispensable for Spo11-DSBs, the absence of either Red1 or Hop1 strongly reduces Spo11-DSBs, notably because Hop1 participates in the recruitment of Mer2, a key Spo11 cofactor for DSB formation (16, 25–32).

After DNA cleavage, Spo11 is released from DNA ends by the Mre11/Rad50/Xrs2 complex in combination with Sae2 but remains covalently attached to an oligonucleotide (33, 34). Spo11-oligo sequencing allowed the establishment of a nucleotide-level resolution Spo11-DSBs genome wide map (10). Once processed, Spo11-DSBs are primarily repaired by homologous recombination with the non-sister homologous chromatids, to eventually yield crossovers (35, 36). This promotes pairing and synapsis of the homologous chromosomes through the synaptonemal complex assembly. The main component of this proteinaceous complex is the transverse filament protein Zip1, and its lateral elements correspond to the chromosomes axes (37). The chromosome axis component Hop1 is eventually removed from the axis upon completion of synaptonemal complex formation, shutting down DSB formation (21, 38–40). Interestingly, the length of the synaptonemal complex covaries with the number of crossovers independently of the genome size and the species (41, 42). In *S. cerevisiae*, the major crossover formation pathway involves the ZMM proteins (Zip1, 2, 3, 4, Spo16, Msh4, 5 and Mer3, (43, 44). Zip3 appears as a specific marker of the resulting “type I” crossovers that show interference (45, 46). Genome-wide distribution of Zip3 colocalizes with Spo11-DSBs and alternates with chromosome axis binding sites (46, 47).

Meiotic homologous recombination is commonly repressed around the loci determining sexual compatibility such as the mating-type loci in fungi, and sometimes this inhibition extends outside of this locus (48). Both the mechanism and the selective advantage of recombination suppression are unclear. Recombination inhibition within the *MAT* locus may prevent loss of heterozygosity at this specific locus and subsequent disequilibrium between the two mating types within the population. In addition, recombination inhibition around the *MAT* locus may ensure linkage between specific alleles that present strong selective advantages when linked (49). As another example of recombination inhibition around the mating-type locus, we previously reported a complete inhibition of meiotic recombination over almost the entire chromosome arm containing the mating type locus of the budding yeast *Lachancea kluyveri* (50). The inhibition spreads over the region of the left arm of *L. kluyveri* chromosome C starting from the left telomere to a point about 10kb to the left of the centromere; we refer to this region as Lakl0C-left. This inhibition of recombination results from the repression of Spo11-DSBs within this ca. 1 Mb long region, corresponding to ca. 8% of *L. kluyveri* genome. So far, the mechanism inhibiting Spo11-DSB formation over such a large genomic region is unknown.

*L. kluyveri* is a *Saccharomycetaceae* yeast that diverged from *S. cerevisiae* prior the whole genome duplication that characterizes the *Saccharomyces* lineage (51). *L. kluyveri* has only eight chromosomes, with a genome size and a gene complement comparable to *S. cerevisiae* (52, 53). As in *S. cerevisiae*, meiosis is efficiently induced by nitrogen starvation and meiotic progression is rather synchronous (50). Interestingly, Lakl0C-left is characterized by a G + C frequency almost 10% higher than the rest of the genome (53). A population genomics survey of *L. kluyveri* led to the proposal that Lakl0C-left arose from an introgression from a yet unknown *Lachancea* species (54). Finally, Lakl0C-left contains almost exclusively early replication origins, in contrast to the rest of the genome (55). Whether any of these Lakl0C-left specificities are at the source of the repression of Spo11-DSBs formation is unknown.

Here, we confirm Spo11-DSB inhibition on Lakl0C-left using a genome wide DSB mapping method (56) and find that the Spo11-DSB hotspot landscape is poorly conserved between orthologous syntenic intergenic regions from *L. kluyveri* and *S. cerevisiae*. Using Hi-C, we showed that *L. kluyveri* chromosomes undergo compaction during meiotic prophase, and that Lakl0C-left compaction is comparable to the rest of the genome, which correlates with a comparable binding of Rec8 as detected by ChIP-seq. By contrast, Hop1 and Red1 are completely absent from Lakl0C-left, likely explaining the lack of Spo11-DSBs. Finally, fluorescence visualization of Zip1, Rec8 and the left and right arms of chromosome C recapitulated these molecular findings and revealed the lack of pairing of Lakl0C-left. Overall, *L. kluyveri* chromosome C appears as a prototypical young sex chromosome.

## Results

### Genome wide Spo11-DSB mapping in *L. kluyveri*

Using pulsed-field gel electrophoresis (PFGE) and Southern blotting, we previously showed that Lakl0C-left is depleted of Spo11-DSBs but flanked by a Spo11-DSB hotspot in the *GPI18* promoter, located about 10 kb to the left of the centromere. We also identified by Southern blot two additional DSB hotspots in the promoters of the *RAS1* and *PIS1* genes that correlate with crossover hotspots (50). To confirm these results with an independent technique and to extend them to the whole genome, we used the Covalent Complexes-sequencing (CC-seq) technique that allows nucleotide-resolution mapping of protein-linked DNA breaks (56).

We performed two independent experiments. After background filtering (methods), we identified a specific signal that clustered into hotspots, as seen for *S. cerevisiae* Spo11-DSBs, and the two replicates showed good reproducibility (Figure 1A and S1). We used the MACS2 peak calling algorithm to define the Spo11-DSB hotspots (57). One replicate showed better signal enrichment than the other (1,933 hotspots vs 1,025 hotspots, Figure S1) and was used for subsequent analyses.

**Figure 1.**
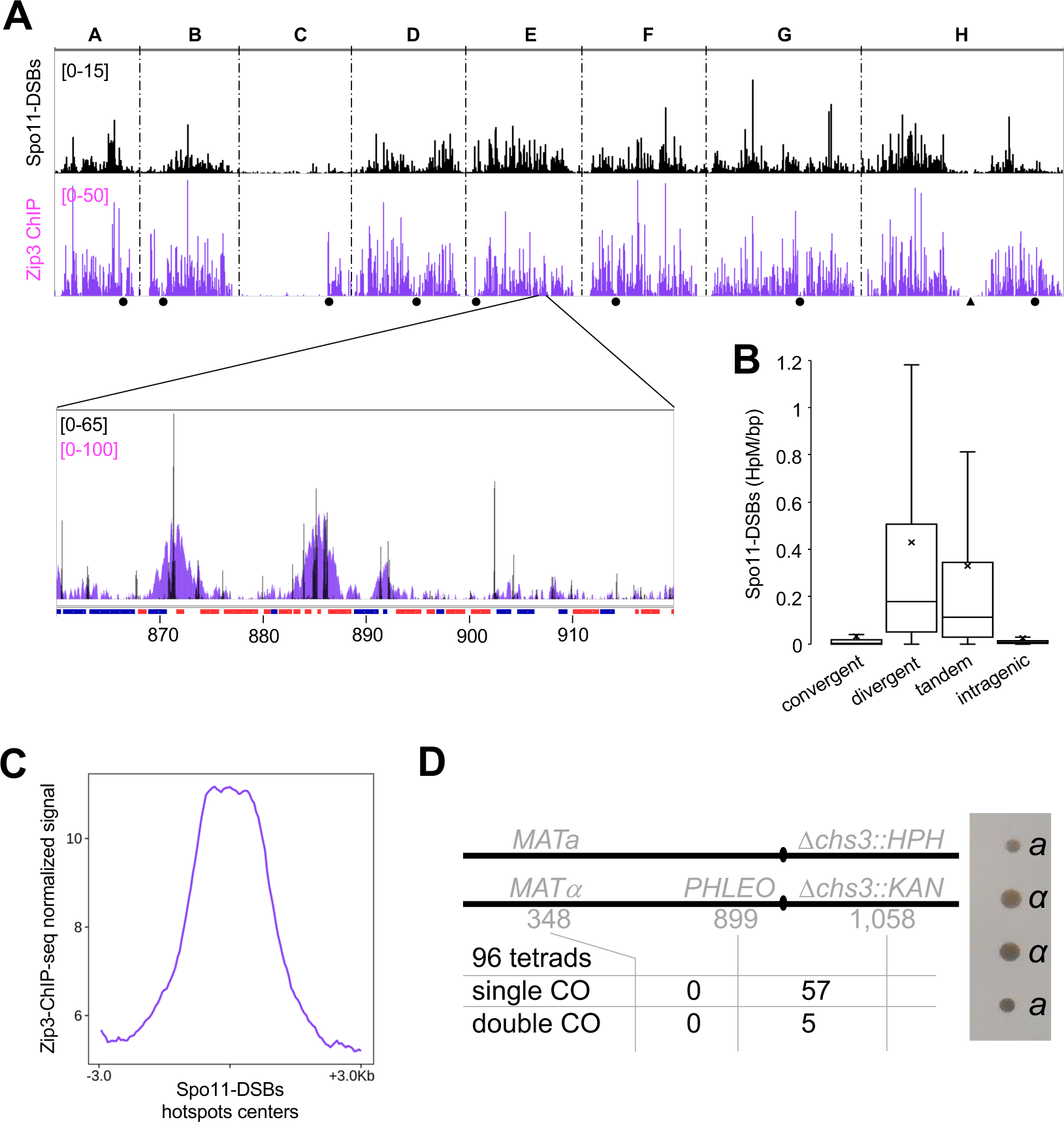
Lakl0C-left is depleted in Spo11-DSBs, Zip3 and crossovers. **A**. Genome-wide CC-seq (Spo11-DSBs, top) and Zip3-FLAG ChIP-seq (bottom) profiles of *L. kluyveri* CBS10367 at six hours post meiosis induction. The eight chromosomes are concatenated in the alphabetical order from A (left) to H (right) and separated by vertical dotted lines. Black discs indicate centromeres. The black triangle indicates the rDNA locus position, which corresponding signal has been masked. The y axis of the CC-seq profile represents the Spo11-DSB count after denoising of a single experiment. The y axis of the Zip3-FLAG ChIP-seq profile represents the mean ChIP-seq signal of five independent experiments. A zoom in a representative region of chromosome E with superimposition of the CC-seq and Zip3-FLAG ChIP-seq signals is shown. Coding regions are represented on the x axis (chromosome coordinates in kilo bases). Transcription is left to right for red boxes and right to left for blue boxes. Maximum values are lower in the genomic view due to visualization constraints. **B**. Box plot representing the distribution of the CC-seq signal according to the type of genomic location. Intergenic regions comprise regions flanked by convergent genes, divergent genes, and genes that are in a head to tail (tandem) configuration. Horizontal bars show the medians and crosses show the means. Boxes delimit the first and third quartiles. The error bars represent minimum and maximum values. **C**. Average signals of five independent Zip3-FLAG ChIP-seq experiments performed 6 hours after meiosis induction piled up around the centers of the 500 strongest Spo11-DSBs hotspots defined by MACS2 from the CC-seq experiment. **D**. Recombination assay in the *L. kluyveri* CBS10367 diploid strain. The two copies of chromosome C are represented as a horizontal black line. Relevant loci are indicated as well as their chromosome coordinates in kb. *PHLEO* was integrated at the Lakl0C-899 locus. The black ovals represent centromeres. The table indicates single and double crossovers (CO) between *MAT* and *PHLEO* and between *PHLEO* and *CHS3* out of 96 tetrads. The picture on the right is representative of a full tetrad with the two *MATa* colonies exhibiting a smaller size than the two *MATα* colonies.

The average DSB hotspot density is 1 per 5.9 kb and the average width was 0.282 kb in *L. kluyveri* compared to 1 per 4.2 kb and 0.409 kb, respectively in *S. cerevisiae* (56). As in *S. cerevisiae*, most DSB signal is in intergenic regions between divergent or tandemly orientated genes (Figure 1B and S2A). Only 13.2% of the DSB signal is within genes and 3.1% is within intergenic regions between convergent genes. In addition, centromere-flanking regions, subtelomeres and the rDNA locus were depleted for Spo11-DSBs compared to the rest of the genome (Figure S3). However, and in sharp contrast with the rest of the genome, a depletion of Spo11-DSBs was observed all along Lakl0C-left, which shows a 12-fold decrease in Spo11-DSBs per base pair compared to the rest of the genome (Figure 1A). Overall, genome-wide mapping of Spo11-DSBs reveals similar properties of Spo11-DSBs distribution between *L. kluyveri* and *S. cerevisiae* chromosomes and confirms the depletion of Spo11-DSBs in Lakl0C-left.

### Spo11-DSBs hotspots are poorly conserved between *L. kluyveri* and *S. cerevisiae*

Spo11-DSBs hotspot positions and strength are remarkably conserved within *Saccharomyces* species, likely because they are mainly found within gene promoters, which are functional genetic elements (12). In *L. kluyveri*, Spo11-DSBs are also mainly located within gene promoters. To test the conservation of Spo11-DSBs hotspot strength between *L. kluyveri* and *S. cerevisiae,* we compared Spo11-DSBs levels in syntenic intergenic regions (sIGRs) *i.e.* intergenic regions flanked by pairs of orthologous genes. Although these two species diverged over 100 million years ago, they share 239 synteny blocks and 2,030 sIGRs (58). The correlation coefficient between the Spo11-DSB hotspots strength of the two species is *r*=0.256 (Figure 2A), showing poor conservation between the two species. Nevertheless, it remains higher than expected by chance since randomization of the corresponding datasets yields no correlation at all (Figure S2B).

**Figure 2.**
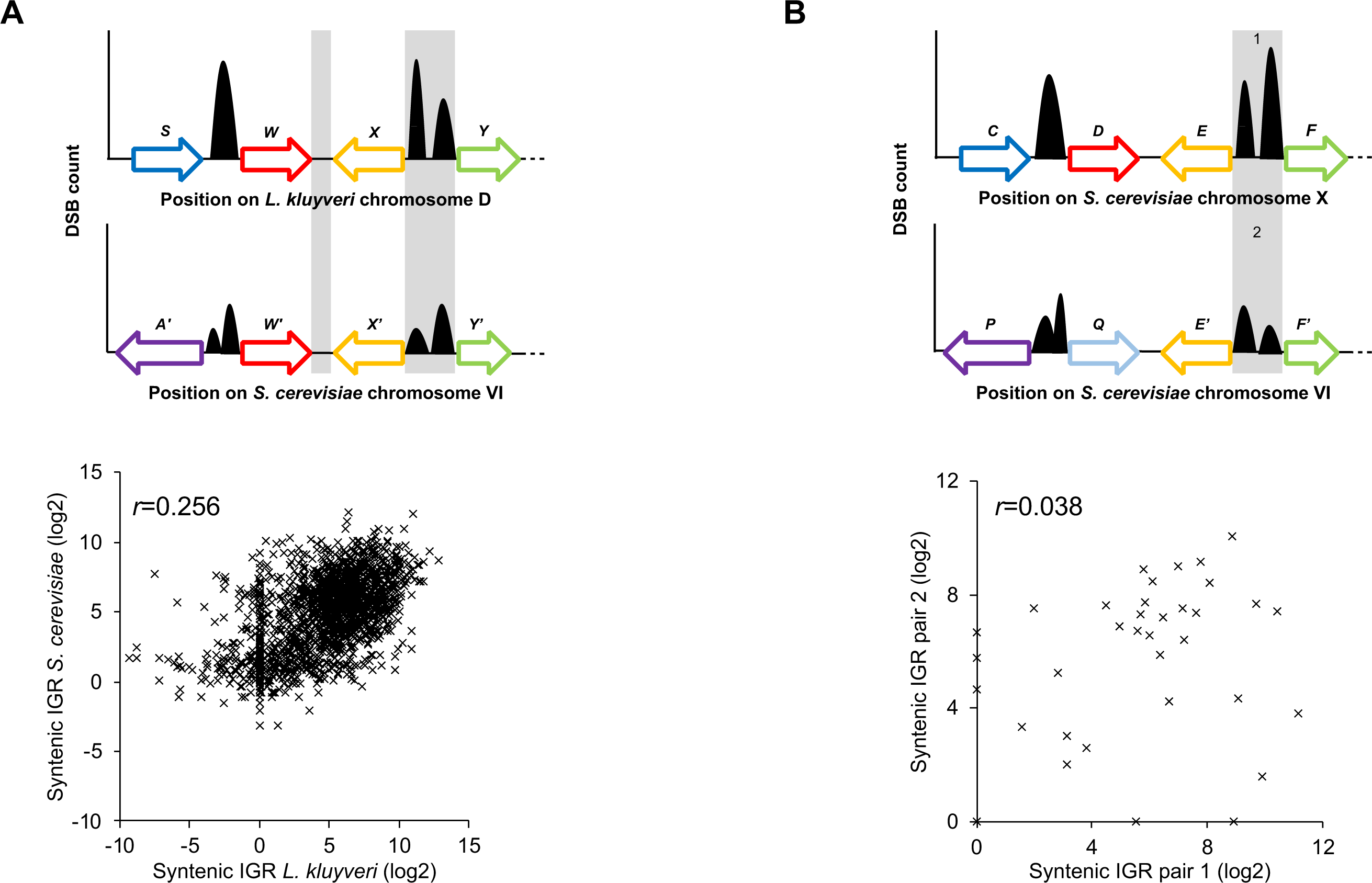
Spo11-DSB signal conservation across syntenic intergenic regions (IGR) between *L. kluyveri* and *S. cerevisiae*, and between *S. cerevisiae* ohnologs. **A.** Top panel represents syntenic intergenic regions (highlighted by grey boxes) in *L. kluyveri* and *S. cerevisiae* that are flanked by orthologous genes couples. The bottom graph shows the comparison of the CC-seq Spo11-DSB signal from 1,956 syntenic IGR between *L. kluyveri* (x axis) and *S. cerevisiae* (y axis) with the correlation coefficient r indicated. **B**. Same as in A except that 39 *S. cerevisiae* intergenic regions flanked by couples of ohnologs are considered.

To assess the conservation of Spo11-DSB hotspot strength on an evolutionary scale broader than that of the *Saccharomyces* species, but independently of *L. kluyveri*, we focused on ohnologs (59). These are the subset of *S. cerevisiae* paralogs that emanated from the whole genome duplication that characterizes the *Saccharomyces* lineage, but that occurred after the divergence between the *Saccharomyces* and the *Lachancea* lineages (60). We identified 39 sIGRs among the 547 ohnolog pairs. Considering the Spo11-oligo data from (10) corresponding to these 39 sIGRs, the correlation coefficient between the Spo11-DSB hotspots strength of these sIGRs is *r*=0.038 (Figure 2B). This shows that, for this subset of intergenic regions, the strength of Spo11-DSBs is not conserved over an evolutionary scale smaller than that of the divergence between the *Lachancea* and the *Saccharomyces* lineages, and therefore suggests different evolutionary constraints on intergenic regions associated with ohnologs versus orthologs.

### Spo11-DSBs sites are enriched in the crossover factor Zip3

In *S. cerevisiae*, the crossover factor Zip3 is enriched around Spo11-DSBs (46). To determine if this is also the case in *L. kluyveri*, we performed ChIP-seq experiments using Zip3 FLAG-tagged on its carboxyl terminus as a bait. As in *S. cerevisiae*, Zip3 is specifically expressed after meiotic induction (Figure S4). We observed Zip3 ChIP-seq enrichment peaks genome wide, with the notable exception of Lakl0C-left and about 250kb left and right of the rDNA locus on Lakl0H (Figure 1A). Piling up Zip3 ChIP-seq signals centered on the 500 strongest Spo11-DSBs hotspots revealed a specific Zip3 enrichment around these Spo11-DSBs hotspots (Figure 1C). Importantly, the absence of Zip3 ChIP-seq signal on Lakl0C-left suggests a complete absence of crossovers in this chromosome arm in the nearly homozygous CBS10367 strain background used here. This population average analysis confirms and generalizes our original observation suggesting, from a limited number of meioses in the artificial NBRC10955 x 67-588 hybrid background, a lack of recombination along this arm (50, 61).

### The lack of meiotic recombination on Lakl0C-left is conserved among *L. kluyveri* isolates and results in a *MAT-*linked growth phenotype

We attributed the complete absence of recombination on Lakl0C-left observed in a hybrid background to the strong depletion of Spo11-DSBs, as measured previously by Southern blot (50) and here by CC-seq in the natural CBS10367 diploid background (54). However, we cannot rule out that a low level of Spo11-DSBs would be enough to induce recombination within Lakl0C-left in this homozygous background, but not in a hybrid background, where sequence polymorphism would prevent recombination through the action of mismatch repair (62–64).

We therefore designed a reporter assay inserting drug resistance cassettes to monitor recombination within two intervals of *L. kluyveri* chromosome C in the nearly homozygous CBS10367 background (Figure 1D). From left to right, the first interval is exclusively on Lakl0C-left and spans about 550 kb from *MAT* to the Lakl0C-899 locus positioned about 110 kb left of the centromere. The second interval starts from Lakl0C-899 to the *CHS3* locus located about 50 kb right of the centromere and is about 160 kb long. The spore viability of the resulting strain is 76% (n=96 tetrads), similar to that of the parental strain (83%, n=39 tetrads). From 96 tetrads with four viable spores, we did not detect a single crossover on the Lakl0C-left 550 kb long interval and we detected 57 single crossovers and five double crossovers on the second 160 kb long interval (Figure 1D). The absence of recombinants on the Lakl0C-left interval in this nearly completely homozygous context shows that inter-homolog recombination is inhibited on Lakl0C-left independently of sequence polymorphism, and that the low Spo11-DSBs level detected does not promote recombination. The absence of meiotic recombination on Lakl0C-left therefore occurs in both the NBRC10955 x 67-588 hybrid and the CBS10367 strains. Given the *L. kluyveri* phylogeny (54), this shows that recombination cessation on Lakl0C-left is ancient and likely common to the entire population.

Remarkably, *MATa* colonies coming from CBS10367 spores were systematically smaller than *MATalpha* colonies (Figure 1D). One explanation could be that there is at least one DNA sequence polymorphism between the *MATa* and *MATα*-associated linkage groups responsible for this sexual dimorphism in the CBS10367 natural diploid. This would agree with the fact that non-recombining regions tend to accumulate heterozygous DNA sequence polymorphisms, as frequently observed for sex chromosomes ((48) and discussion). Alternatively, differentially expressed genes between *MATa* and *MATalpha* cells could be responsible for this growth phenotype, although this is not the case in other species including *S. cerevisiae*.

### Hi-C reveals similar compaction level of all *L. kluyveri* meiotic chromosomes

We wondered if the lack of recombination on Lakl0C-left could result from a peculiar chromosome structure that would differ from the canonical compaction of chromosomes during meiosis prophase into cohesin-mediated chromatin loops (14). Using Hi-C (Lieberman-Aiden et al. 2009; Muller et al. 2018), we generated genome-wide contact maps of eight *L. kluyveri* meiotic timepoints harvested between zero- and eight-hours after meiosis induction. ∼70% of cells passed the second meiotic division at the eight-hours timepoint (Figure S5A). Overall, the contact maps of *L. kluyveri* meiotic Hi-C resemble those from *S. cerevisiae* (Figure 3A) (15, 17). Centromere clustering is clearly visible at t=0 (Rabl configuration) and is progressively lost (Figure 3A, S5B). Chromosomes gradually individualize, as shown by the increase and decrease of intra- and inter-chromosomal contacts, respectively. Compaction was maximal five hours following entry into meiosis (Figure S5C). Enrichment in intrachromosomal contacts involves pairs of loci within the 10 to 200 kb range, as illustrated both by the thickening of the main diagonals on the normalized contact maps (Fig. 3 A and B) and the shoulder made by the distance law, reflecting the genomic average contact frequencies over increasing genomic distances (Figure 3C). This compaction most likely results from the folding of chromatin into loops by the cohesin complex (15, 17, 18). Remarkably, the Hi-C pattern displayed by the Lakl0C-left was comparable to the other chromosomes. The intrachromosomal contact frequency and the distance law of the Lakl0C-left arm displayed nearly the same compaction throughout the meiotic time course than the other chromosome arms of similar lengths (Figure 3C, S5D and E). In addition, based on similar interchromosomal contact frequencies compared to the other chromosome arms, Lak0C-left is not isolated from the other chromosomes (Figure 3A, S5C and D). Overall, Lakl0C-left is folded into cohesin mediated loops, whose scores, length and kinetic of formation are similar to loops analyzed on other chromosomal arms (Figures 3B, D and E, S5E and F). In conclusion, this shows that the lack of recombination of Lakl0C-left is not associated with any specific structure detectable by Hi-C, and that Lakl0C-left undergoes similar compaction by cohesins as all other chromosomes.

**Figure 3.**
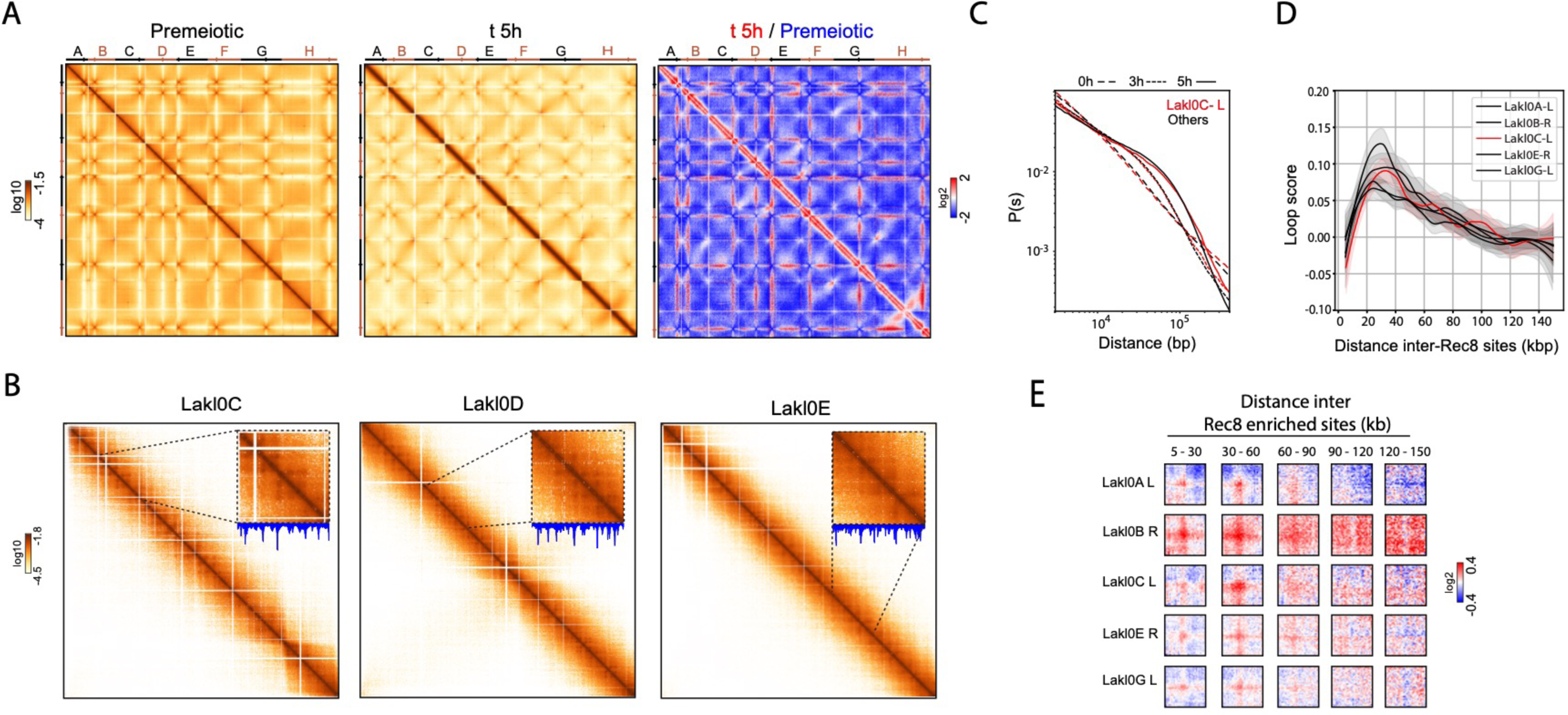
Lakl0C-left compaction along meiotic prophase is similar to other chromosomal arms. **A.** Contact maps of the whole genome of *L. kluyveri* before (left) and five hours (middle) after induction of meiosis. Ratio maps of five hours over premeiotic chromosomes (right). Bin size, 10 kb. **B.** Contact maps of chromosome C (left), D (middle) and E (right) at five hours after meiosis induction. Bin size, 1 kb. Magnified insets show portions of 200 kb, with the corresponding Rec8 ChIP over untagged ratio profiles, determined at four hours post meiosis induction. **C.** Distance laws of Lakl0C-left compared to the mean of ∼1Mb of different chromosome arms before, three hours and five hours post meiosis induction. **D.** Spectrum of loop score detected at five hours post meiosis induction at the intersection of Rec8 peaks detected at four hours post meiosis induction, as a function of the distance separating the pair of Rec8 peaks (kb). **E.** Aggregated ratio maps of 29 kb windows (five hours post meiosis induction) centered on pairs of Rec8 peaks (four hours post meiosis induction) over randomly chosen positions in the same inter-distance range. Bin size: 1 kb.

### Meiotic axis proteins Hop1 and Red1 are absent from Lakl0C-left

The compaction of Lakl0C-left suggests the loading of cohesins, including the meiosis specific Rec8 kleisin subunit. To assess this, we performed ChIP-seq analysis of Rec8, FLAG-tagged at its carboxy-terminus, which was specifically expressed after meiosis induction and show retarded forms on western blots, as in *S. cerevisiae* (Figure S4) (Brar et al. 2006). *L. kluyveri* Rec8 ChIP-seq profiles are similar to those of *S. cerevisiae* (Figure 4). The Rec8 ChIP-seq signal first appears early (1h) after induction of meiosis around all centromeres (Figure S6) (32). Four hours after meiosis induction, the Rec8 ChIP-seq signal is then distributed in peaks throughout the genome and accumulates in convergent intergenic regions, likely as a result of transcription (Figures 4, S6, S7A and B) (23, 66). In addition, Rec8 ChIP-seq peaks alternate with the peaks of Zip3 ChIP-seq and Spo11-DSBs CC-seq (Figure S7A and C). Such a distribution is compatible with the loop-axis structure of meiotic chromosomes, with Spo11-DSBs formed within the loops at the basis of which lie the axis proteins including Rec8 (16). Importantly, the loci at the basis of such chromatin loops detected as discrete dots away from the diagonal in the Hi-C experiments are enriched in Rec8 (Figures 3B and E) (67). Furthermore, Rec8 ChIP-seq signal is decreased in subtelomeric regions (Figure S8). We noted a slight reduction of Rec8 ChIP-seq signal on Lakl0C-left relative to the rest of the genome (Figure S7B, compare right and left panels, respectively). However, it comes from a nonspecific sequencing bias, also observed in the untagged control, that is likely due to the elevated GC content of Lakl0C-left (68). Considering this sequencing bias, the overall Lakl0C-left Rec8 ChIP-seq signal is comparable to the rest of the genome, which is in agreement with the Hi-C results showing comparable compactions.

**Figure 4.**
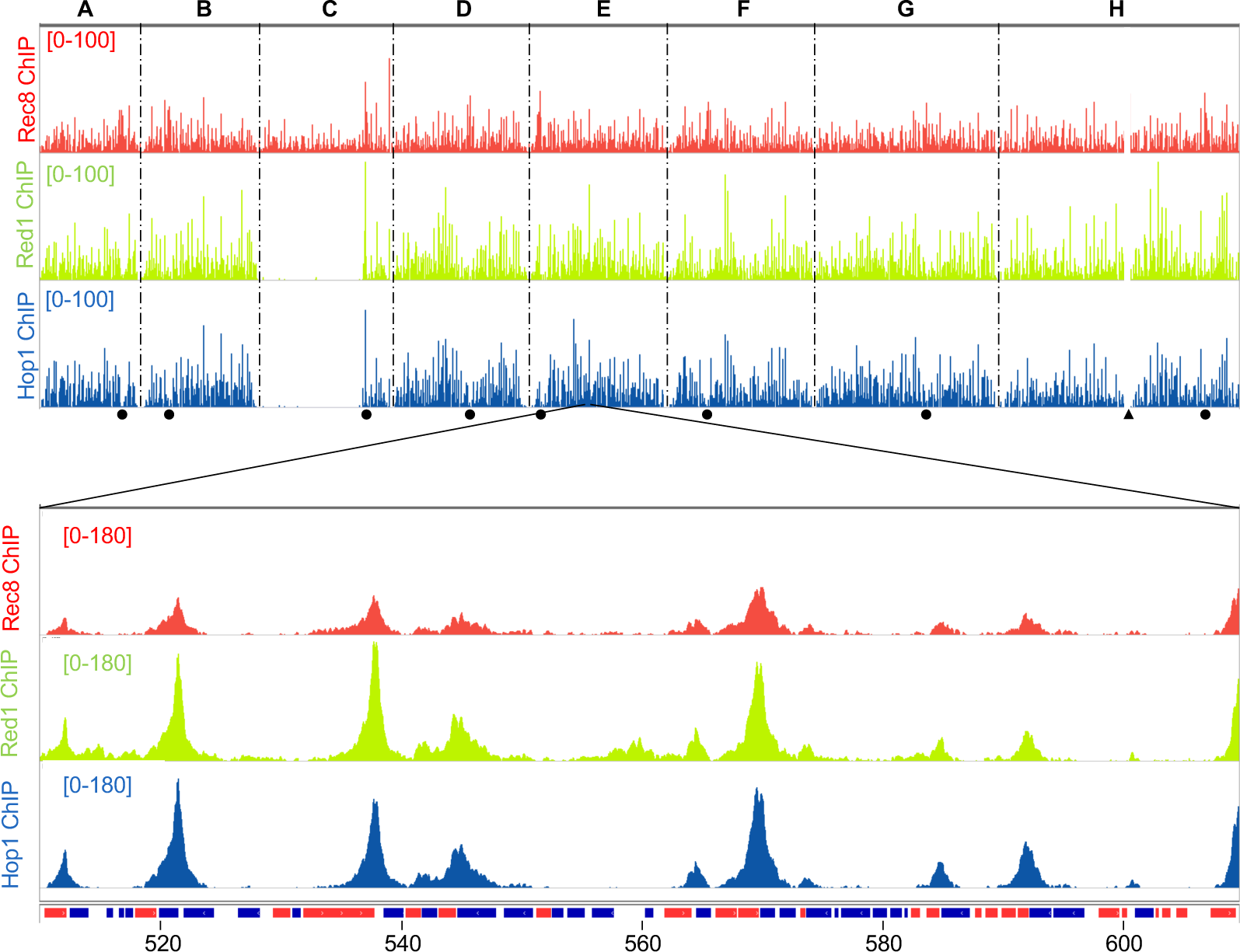
ChIP-seq of chromosome axis proteins Rec8-FLAG, Hop1-FLAG and Red1-FLAG. The top panel represents the genome-wide ChIP-seq signals from two independent experiments six hours after meiosis induction for the three proteins. Same legend as in Figure 1. The bottom panel is a close-up view of a representative genomic region of chromosome E illustrating the strong similarity between the ChIP-seq profiles of the three proteins. Genes coding regions are represented as in Figure 1A.

Coalescence of cohesins participates in the formation of the chromosome axes that also comprise Hop1 and Red1, needed for Spo11-DSBs formation (26–28, 30, 31). Since Rec8 loads normally on Lakl0C-left, we investigated if Red1 and Hop1 were also loaded. Like for Rec8, we tagged Red1 and Hop1 with FLAG at their carboxy-termini. As in *S. cerevisiae*, both proteins are specifically expressed after meiosis induction and show retarded forms on western blots (Figure S4) (69, 70). In addition, Red1 and Hop1 ChIP-seq signals form peaks that strongly colocalize with Rec8 ChIP-seq peaks (Figures 4, S6, S7A and D). However, several cases of Rec8 peaks without associated Hop1 or Red1 peaks were noted. First, Red1 and Hop1 are depleted around centromeres, unlike Rec8 (Figure S8). This depletion of Red1 and Hop1 coincides with the depletion of the Spo11-DSB signal (Figures 1A, S3 and S7A) and the decrease in recombination (50), as seen in *S. cerevisiae*. Second, around the rDNA locus, while neither Rec8 nor Red1 ChIP-seq signal shows any decrease compared to the rest of the genome, the Hop1 ChIP-seq signal is decreased (Figure S8, chromosome H). Strikingly, this decrease occurs on a much narrower region than that of Zip3. Finally, the most remarkable discrepancy between Rec8 and Hop1/Red1 signal was observed on Lakl0C-left, where virtually no Red1 or Hop1 ChIP-seq signal was observed despite a normal Rec8 loading (Figure 4). While in *S. cerevisiae* Hop1- and Red1-rich or poor regions have been described (16, 71), this is the first time, to our knowledge, that a complete depletion over an almost entire chromosome arm is observed. Given the importance of Hop1 and Red1 for Spo11-DSBs formation, it is tempting to speculate that their absence is directly responsible for the absence of Spo11-DSBs formation on Lakl0C-left.

### Spo11-DSBs formation inhibition on Lakl0C-left is independent of Tel1

In the absence of Tel1, Spo11-DSBs increase in *S. cerevisiae*, although to a lesser extent than what is observed in the absence of its ATM homolog in mice, notably on the non-homologous parts of the sex chromosomes (72–74). We therefore tested by PFGE followed by Southern blot meiotic Spo11-DSBs formation in the *tel1* null mutant. The overall Spo11-DSB profile is similar in the reference and *tel1* backgrounds, notably on chromosome C (Figure S9). This shows that Spo11-DSB depletion on Lakl0C-left is independent of Tel1.

### The lack of recombination on Lakl0C-left is independent of its early replication timing

Spo11-DSB formation is coordinated with meiotic DNA replication and occurs on fully replicated chromosomes (75–77). Interestingly, Lakl0C-left is replicated earlier than the rest of the genome due to the prevalence of early firing replication origins in vegetative cells (55, 78). Although studies in *S. cerevisiae* suggest that this would result in early recruitment of DSB factors and early DSB formation on Lakl0C-left (27, 77), we wondered whether, in *L. kluyveri*, early replication timing would prevent Spo11-DSBs formation, and hence recombination. Therefore, we subdivided the genome in four replication quartiles of eight minutes each based on the known vegetative replication timing (55, 78) and assessed Spo11-DSBs and Zip3-ChIP-seq signals. This partitioning shows that the early replication quartile encompasses a smaller genomic fraction than the mid-early and the late replication quartiles, but a larger genomic fraction than the late replication quartile (Figure 5A). In addition, it shows that Lakl0C-left contains mainly early and mid-early replicated regions. Considering the genomic fraction encompassed by each quartile, it appears that both the early and mid-early replication quartiles contain a significant fraction of the Spo11-DSBs and the Zip3-ChIP-seq signals, which vary less than two-fold between the four quartiles (Figure 5B and C). This shows that the early and mid-early replicated regions from *L. kluyveri* undergo Spo11-DSBs and recombination to a comparable extent as the rest of the genome. Hence, the depletion in Spo11-DSBs and subsequent recombination on Lakl0C-left is not related to its early / mid-early replication timing. Further regulation therefore exists to explain the Spo11-DSBs depletion on Lakl0C-left. Interestingly, the earliest quartile shows slightly fewer Spo11-DSBs than the next two quartiles while it shows slightly more Zip3-ChIP-seq signal, suggesting potential differences in the outcome of early versus late Spo11-DSBs.

**Figure 5.**
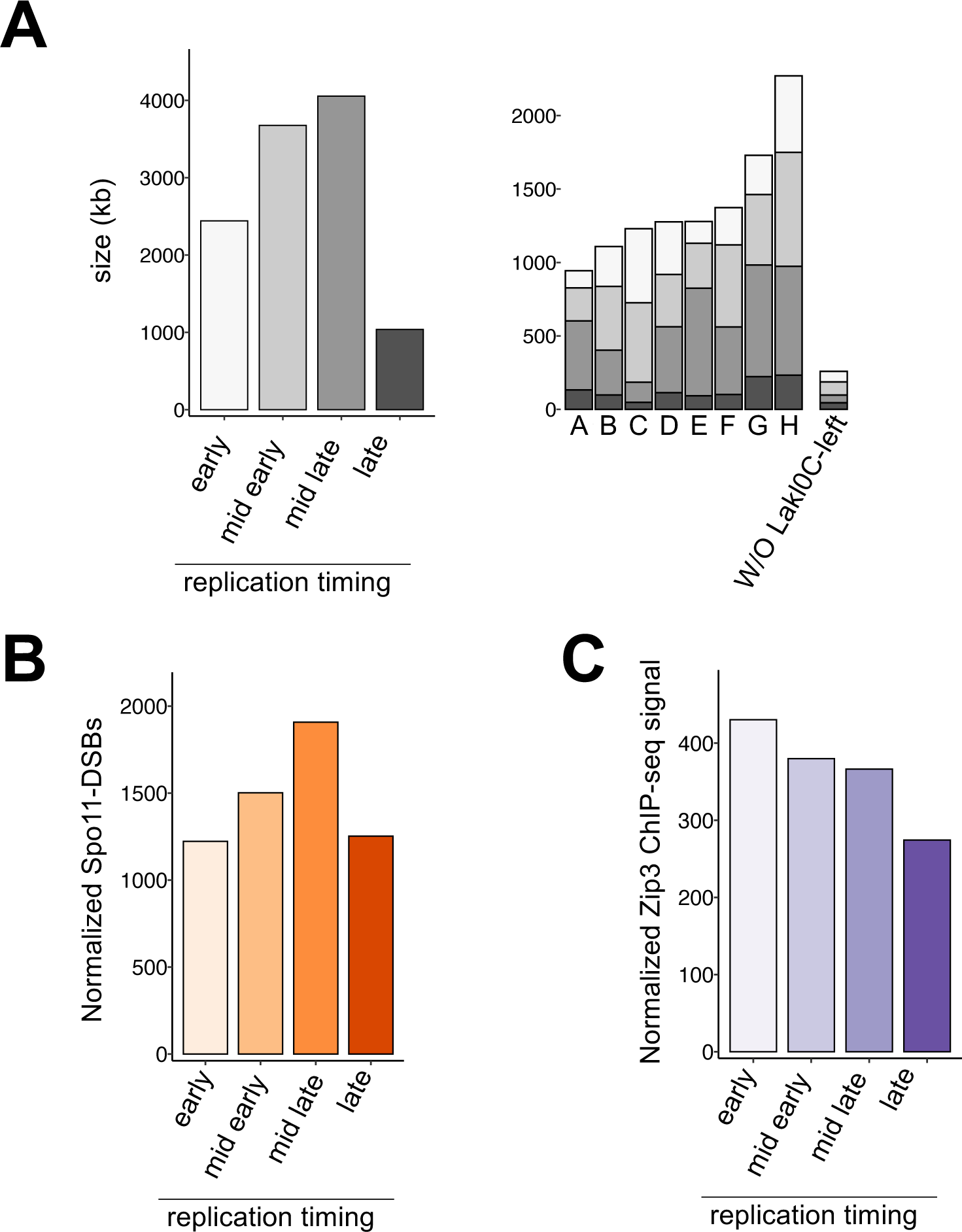
Spo11-DSBs and Zip3-FLAG ChIP-seq enrichment as a function of the replication timing. **A**. *L. kluyveri* genome was partitioned in four groups encompassing the entire vegetative replication phase. Each group corresponds to a period of eight minutes. The left graph represents the fraction of the genome (in kb) encompassed by each replication group. The right graph represents the fraction of each replication group within each chromosome. W/O Lakl0C-left: region of *L. kluyveri* chromosome C excluding Lakl0C-left. **B**. Fraction of the Spo11-DSBs signal at six hours post meiosis induction included in each replication group normalized by the genomic fraction of each group. **C**. Fraction of the Zip3-FLAG ChIP-seq signal at six hours post meiosis induction included in each replication group normalized by the genomic fraction of each group.

### Lakl0C-left does not synapse

Recombination is required to initiate synaptonemal complex formation and homolog synapsis in *S. cerevisiae* (44). Based on this, the absence of recombination on Lakl0C-left may prevent synapsis in this chromosome arm, unless synaptonemal complex formation and synapsis could spread through its entire length after initiating either right of it or at the very left telomere from which recombination cannot be genetically mapped. To test this prediction, we performed immunofluorescence staining of *L. kluyveri* meiotic chromosome spreads using a *S. cerevisiae* anti-Zip1 antibody that cross reacts in *L. kluyveri* and an anti-FLAG antibody directed against a FLAG tagged version of Rec8. Experiments were performed in the absence of Ndt80 to block cells at the pachytene stage. As expected, we observed continuous Zip1 staining of *L. kluyveri* meiotic chromosomes (Figure 6A and B). We observed on average nine Zip1 continuous lines per nucleus in agreement with the chromosome number. Interestingly, the cumulated length of Zip1 lines in *L. kluyveri* is 13.86 micrometers on average compared to 28 micrometers in *S. cerevisiae* (Figure 6C). This ca. two-fold longer meiotic chromosome size agrees with the ca. four-fold higher recombination frequency in *S. cerevisiae* compared to *L. kluyveri* (50). It also suggests longer loops in *L. kluyveri* than in *S. cerevisiae*. Measurement of the width of the DAPI signal of pachytene chromosomes spread on the same slide showed this width was on average 1.26 micrometers in *L. kluyveri* and 0.83 micrometers in *S. cerevisiae*, therefore supporting longer chromatin loops in *L. kluyveri* (Figure 6D and E).

**Figure 6.**
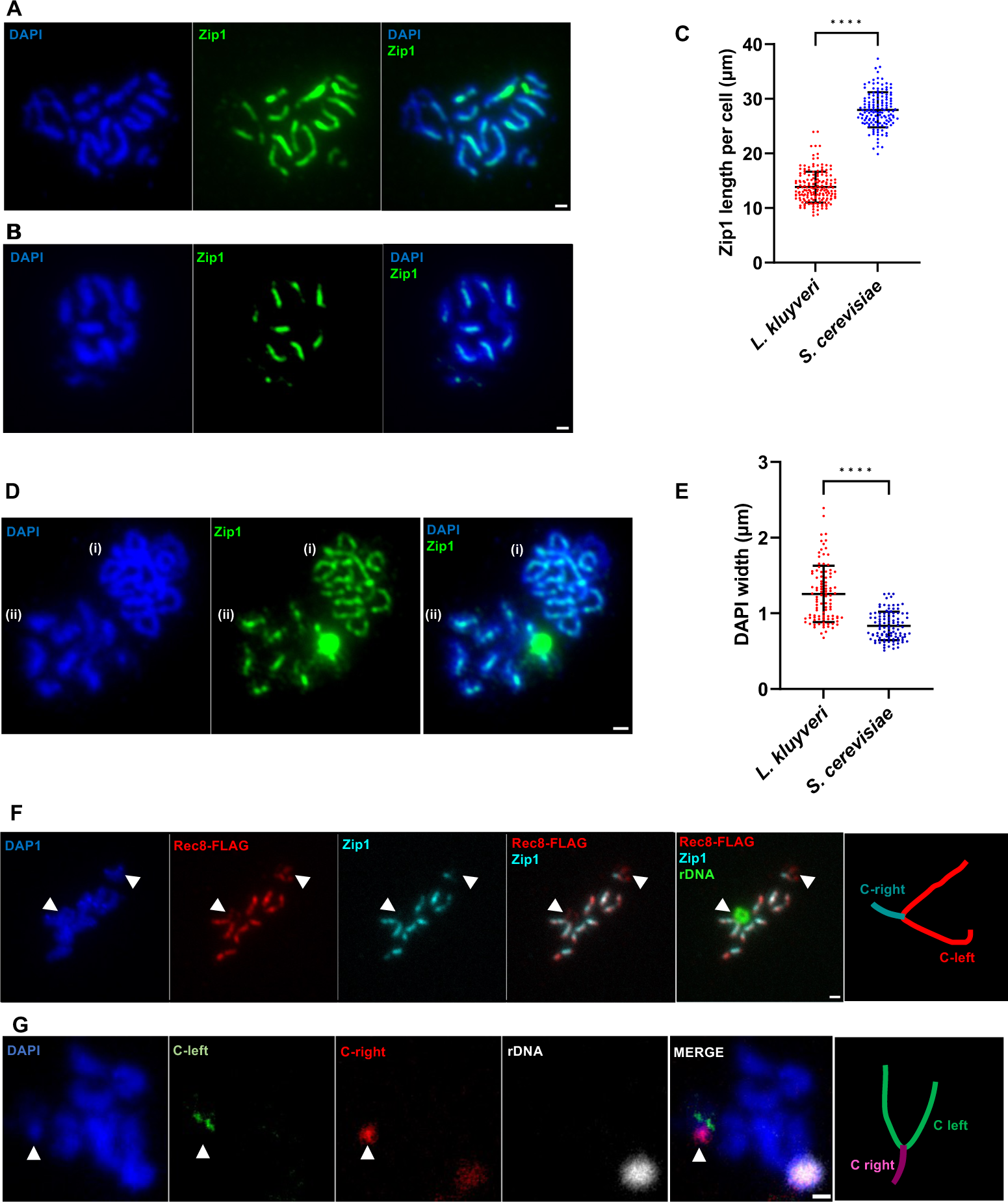
Cytology of *L. kluyveri* meiotic chromosomes. **A**. Pachytene spread nuclei of *S. cerevisiae ndt80* mutant strain (BLY1808) 12h after sporulation induction stained with DAPI (blue) and anti-Zip1 antibody (green). **B.** Pachytene spread nuclei of *L. kluyveri ndt80* mutant strain (LAKL220) 13.5h after sporulation induction stained with DAPI (blue) and anti-Zip1 antibody (green). **C.** Quantification of the total length of Zip1 lines per cell in μm. Black lines show means ± SD. *L. kluyveri:* 13.86 ± 2.81 μm (n= 170), *S. cerevisiae:* 28 ± 3.2 μm (n= 139). The average number of Zip1 lines observed and quantified per cell is 8.7 ± 0.76 for *L. kluyveri* and 14.24 ± 1.32 for *S. cerevisiae.* **D.** Pachytene spread nuclei of (i) *S. cerevisiae ndt80* mutant strain (BLY1808) 12h after sporulation induction and (ii) *L. kluyveri ndt80* mutant strain (LAKL220) 13.5h after sporulation induction stained with DAPI (blue) and Zip1 (green). *S. cerevisiae* and *L. kluyveri* spread nuclei were mixed on the same slide to make sure they undergo the same treatments. **E.** Quantification of DAPI signal width from panel D which reflects the chromosome loops size. Black lines show mean ± SD. *L. kluyveri:* 1.26 ± 0.37 μm (n= 100), *S. cerevisiae:* 0.83 ± 0.19 μm (n= 104). In panel C and E, p< 0.0001, Mann-Whitney U test. **F and G**. Pachytene spread nuclei of *L. kluyveri ndt80* mutant strain (LAKL227) 13.5h after sporulation induction. In F, staining was performed with DAPI (blue), anti-FLAG antibody against Rec8-FLAG (red), anti-Zip1 antibody (cyan) and an rDNA FISH probe (green). The left arrowhead points at the rDNA locus. The right arrowhead points at the inferred chromosome C right unsynapsed arms. In G, staining was performed with DAPI (blue), Lakl0C-left FISH probe (green), Lakl0C-right FISH probe (red) and an rDNA FISH probe (white). The arrowhead points at chromosome C. The structure of chromosome C inferred from staining experiments is drawn (not to scale) on the right of panels F and G. Scale bars = 1 µm.

Rec8 staining colocalized with Zip1 staining except at two regions that exhibited Rec8-only staining (Figure 6F). One of these regions exhibit a circular structure emanating from a chromosome which corresponds to the rDNA locus as demonstrated by staining with a specific FISH probe. The second region consists of a “V” shape emanating from a base stained by both Zip1 and Rec8. This overall “Y” shape chromosome could be compatible with chromosome C, with the base of the Y corresponding to the synapsed right arm, and the arms of the Y corresponding to the left arms entirely unsynapsed. Corroborating this hypothesis, using FISH probes specific of the left and the right arms of chromosome C, we could detect that the FISH probe specific of the right arm of chromosome C stains a DAPI dense region, while the FISH probe specific of the left arm of chromosome C stains two nearby areas less DAPI dense (Figure 6G). Such signals support synapsed right arms of chromosome C and unsynapsed left arms. Consistently, at five hours post meiosis induction, the distance law of Lakl0C-left shows a higher probability of contact at longer range (> 100 kb) than other chromosome arms, which could reflect the ability of the unsynapsed arm to contact distant regions (Figure 3C). In agreement with the ChIP-seq experiment that revealed the presence of Rec8 along Lakl0C-left and the Hi-C experiment that revealed normal compaction of Lakl0C-left, the Y structure is also entirely stained by Rec8. Finally, the fact that the arms of the Y structure are separated further suggests that there is no recombination within the left telomere of chromosome C in regions that could not be genetically mapped nor assessed by our ChIP-seq or CC-seq approaches. Overall, our analysis of *L. kluyveri* meiotic chromosomes spreads reveals that Lakl0C-left does not synapse. This shows that recombination and subsequent synapsis initiated outside Lakl0C-left is not enough to spread through this entire region. Recombination seems therefore also required in *L. kluyveri* to locally initiate homolog synapsis, as in *S. cerevisiae*.

## Discussion

The lack of meiotic recombination on Lakl0C-left implies an absolute genetic linkage and an absence of sequence homozygosis by gene conversion during meiosis. This chromosome arm that contains the *MAT* locus therefore relies exclusively on mitotic recombination to break genetic linkages and homogenize mutations. It is therefore expected that Lakl0C-left genes would exhibit a lower purifying selection than the other genes of the genome. This is actually the case, since Lakl0C-left genes show a higher ratio of non-synonymous (dN) to synonymous (dS) substitution rates (dN/dS) than the rest of the genome (79). It is also expected to find a higher density of heterozygous sequence polymorphisms on Lakl0C-left than on the rest of the genome, with at least one possibly being responsible for the different growth phenotypes of *MATa* and *MATα* spores from the CBS10367 isolate. In agreement with this expectation, analysis of the CBS10367 diploid genome with respect to the CBS3082 reference genome revealed a significantly higher fraction of heterozygous SNPs and indels on Lakl0C-left compared to the other chromosomes excluding the highly variable subtelomeric regions, with 1.53 vs 0.11 heterozygous SNPs per kb and 0.33 vs 0.018 heterozygous indels per kb, respectively (Table 1 and Figure S10A). Despite this higher SNP density, the most telomeric ca. 200 kb left of chromosome C are almost completely devoid of heterozygous SNPs, potentially reflecting homozygosis by mitotic recombination (G2 crossover or break-induced replication) in the recent history of the strain. Finally, the density of homozygous sequence polymorphisms on Lakl0C-left existing between CBS10367 and CBS3082 is comparable to the rest of the genome, with 25.29 vs 23.42 homozygous SNPs per kb, respectively, and is much higher than the density of heterozygous sequence polymorphisms mentioned above (Figure S10B). Given the absence of meiotic recombination on Lakl0C-left that is likely common to all *Lachancea kluyveri* isolates, this suggests a leading role of mitotic recombination in yielding homozygous sequence polymorphisms in *L. kluyveri*.

**Table 1.**
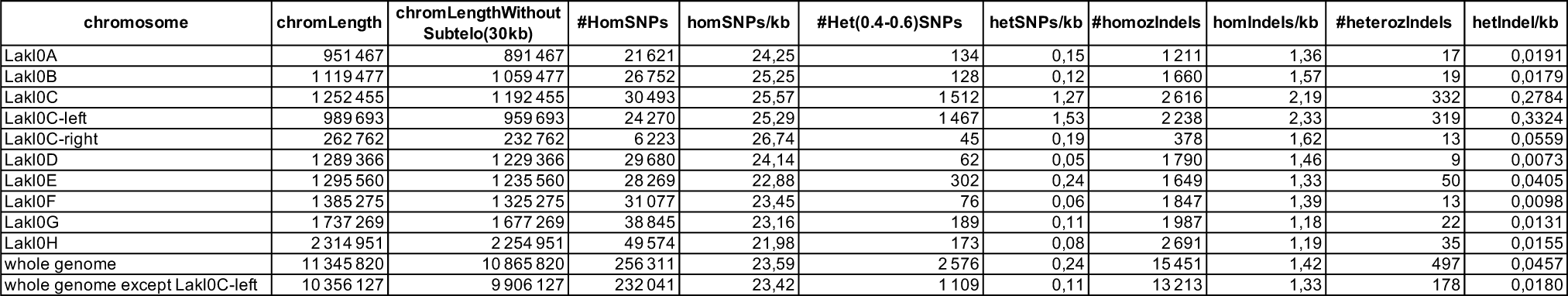

Having determined the Spo11-DSBs landscape of *L. kluyveri*, we could assess the conservation of the strength of Spo11-DSBs hotspots between *L. kluyveri* and the reference *S. cerevisiae* by comparing the frequency of Spo11-DSBs within syntenic intergenic regions as in (12). Unlike within the *Saccharomyces* species, the strength of Spo11-DSBs hotspots is only poorly conserved between *L. kluyveri* and *S. cerevisiae*, and does not result in conserved crossover hotspots (50). We figured out that the ohnologs provide another relevant dataset to look at the conservation of Spo11-DSBs hotspots within a single genomic environment at a broad evolutionary scale, larger than that of the *Saccharomyces* clade but shorter than that separating *S. cerevisiae* and *L. kluyveri*. Although the sample size is small, the strength of the Spo11-DSBs signal within syntenic inter-ohnolog regions is not conserved at all. In both cases it is not possible to distinguish between hotspot erosion, already visible within the *Saccharomyces* clade, and different chromosomal environments or selective constraints as a source of this lack of Spo11-DSBs hotspot strength conservation. However, this shows that the strength of Spo11-DSBs hotspots that do not rely on sequence specific elements tend to be not conserved at such an evolutionary scale within budding yeasts, which is about 100 million years (80).

In most species studied so far, the number of meiotic crossovers per chromosome is between one and three. *S. cerevisiae* and *S. pombe* are among the outliers with much more crossovers per chromosome (81, 82). With a comparable genome size and twice fewer chromosomes, *L. kluyveri* has ca. four-fold fewer crossovers per meiosis than *S. cerevisiae* (50). *L. kluyveri* therefore resembles most species, with the exception of Lakl0C-left that likely involves an additional mechanism preventing recombination. We observed a ca. two-fold longer synaptonemal complex length in *S. cerevisiae* and a shorter chromosome width as measured by DAPI staining. Given the evolutionary conservation of the dimensions of the “loop-base module” (42) and the similar genome size of *S. cerevisiae* and *L. kluyveri*, this suggests less but longer loops in *L. kluyveri* compared to *S. cerevisiae*. The correlation of chromosome axis length and frequency of crossovers between these two closely related species is reminiscent of the covariation of the frequency of crossovers per nucleus with the chromosome axis length (41, 83). While this latter point suggests a constant crossover frequency per chromatin loop within a given species, the actual crossover frequency per chromatin loop might be different between *L. kluyveri* and *S. cerevisiae.* Overall, given the strong difference in terms of recombination frequency and chromosome axis length between *S. cerevisiae* and *L. kluyveri*, the detailed comparison of their meiotic chromatin loops and associated genes and proteins may be a way to understand what governs the meiotic recombination frequency of a given species. Notably, the Pds5 cohesion maintenance factor and the NuA4 histone acetyltransferase complex catalytic subunit Esa1 were shown to contribute significantly but independently to meiotic chromosome axis length (84, 85). It will be interesting to determine their contributions in the difference in meiotic chromosome axis length between *L. kluyveri* and *S. cerevisiae*.

The Y structure of Lakl0C chromosome during pachytene, with only the base of the Y stained by Zip1, suggests that the synaptonemal complex does not form along Lakl0C-left, although it is present on the right arm of this chromosome. While recombination is essential to initiate synaptonemal complex formation in *S. cerevisiae* and likely in *L. kluyveri* (44), our results suggest that either the extent of polymerization of the synaptonemal complex in *L. kluyveri* is limited and requires initiation points not too far apart to cover chromosomes entirely, and/or that something is specifically inhibiting synaptonemal complex polymerization on Lakl0C-left after its priming outside of Lakl0C-left. Alternatively, the failure to polymerize could be due to the absence of pre-existing Red1 or Hop1, as discussed below.

Our most striking finding is the complete absence of the meiotic axis proteins Hop1 and Red1 from Lakl0C-left. The abundance of these proteins is known to vary along chromosomes. Hop1 is notably enriched in regions with higher recombination activity (16). However, the complete absence of Hop1 and Red1 from a chromosome region that encompasses ca. 8% of the entire genome has not been reported so far, to our knowledge. Despite their phylogenetic divergence, the regulation of meiotic recombination is highly similar between *S. cerevisiae* and *L. kluyveri*. Most if not all relevant meiotic recombination genes are conserved between the two species. This contrasts with all the other *Lachancea* species that lost most genes from the ZMM pathway right after the divergence of *L. kluyveri* from the rest of the clade (58, 86). Notably, Hop1, Red1 and Rec8 display similar properties between *S. cerevisiae* and *L. kluyveri*. These include the early recruitment of Rec8 around centromeres and the subsequent recruitment of Rec8 in discrete peaks between convergent genes throughout the genome that strongly colocalize with both Hop1 and Red1 peaks. This suggests conserved recruitment regulation of these proteins. In *S. cerevisiae*, the main Hop1 and Red1 recruitment pathway is through Rec8 that first recruits Red1 which subsequently recruits Hop1 (16, 23). A second recruitment pathway independent of Rec8 and relying on the PHD domain of Hop1 was recently reported (24). The complete absence of both Hop1 and Red1 from Lakl0C-left suggests that these two pathways are inhibited in this entire 1 Mb long region. Whether this relies on the presence of an inhibitory factor, or the absence of an activating factor is so far unknown. Although the specific early replicating profile was a plausible explanation for this chromosome-wide behavior, we showed that other early replication regions are not significantly depleted in Spo11-DSBs and Zip3 ChIP-seq signals. Another hypothesis is that this behavior is related to the ca. 12% higher GC content of Lakl0C-left than the rest of the genome, *i.e.* 52.9 vs 40.4% respectively (53, 87). Interestingly, Heldrich et al (24) showed that the direct recruitment of Hop1 to genomic islands relied on the local DNA sequence. Therefore, one possibility is that Lakl0C-left sequence is incompatible with direct Hop1 recruitment, for reasons that remain to be determined. However, Rec8 is properly loaded on Lakl0C-left as detected by ChIP-seq where it ensures proper chromosome condensation as detected by Hi-C and directly by Rec8 immuno-staining. Hence, while the unusual composition bias of Lakl0C-left could explain the absence of direct recruitment of Hop1, it likely does not explain the inhibition of Hop1 recruitment by Rec8.

Extensive recombination suppression is seen on sex chromosomes, self-incompatibility loci in plants and mating type loci in fungi and algae (48). In the case of fungi, the rationale for the initial recombination suppression at the mating type loci is probably to avoid breaking linkage between genes controlling correct mating. In the budding yeasts *S. cerevisiae* and *L. kluyveri*, such genes include *MAT ALPHA1* and *ALPHA2* genes in the Matα genotype, and *MATA1* and *MATA2* in the Mata genotype, although no known role of *MATA2* has been determined so far. Interestingly, recombination cessation extends beyond the *MAT* locus in many fungi, but the selective advantage is not always clear (48). In the case of *L. kluyveri*, the recombination cessation is likely concomitant with the introgression of Lakl0C-left, therefore relatively recent and includes only one evolutionary stratum covering 80% of chromosome C. No such recombination inhibition is observed in neighbor species including *L. waltii* and *S. cerevisiae*. Interestingly, although meiotic recombination is inhibited, the accumulation of homozygous sequence polymorphisms between isolates suggests that mitotic recombination is taking place equally in Lakl0C-left and the rest of the genome, and that it plays the major role in *L. kluyveri* genome evolution. Overall, while no clear selective advantage has been identified so far for the recombination cessation on Lakl0C-left, the mechanism reported here, *i.e*. the lack of recruitment of meiotic chromosome axis proteins, could be responsible for the extension of recombination cessation around mating type loci in many fungal genomes and play an important role in the emergence of sex chromosomes that rely on the cessation of meiotic recombination (48, 88).

## Methods

### Strains and media

All yeast strains used in this study are derivatives of *L. kluyveri* CBS10367 natural diploid isolate used for its efficient meiotic entry and progression (54, 79). Strain genotypes are listed in Table S1 as well as plasmids used. Gene editing was performed by PCR-mediated gene targeting followed by PCR analysis. Long homology fragments (ca 500 bp long) were used because short homology fragments are not efficient for gene targeting in *L. kluyveri*. Combining long homology fragments with selective markers was made by overlapping PCR or Gibbson cloning (Cat# E5520S, NEB). Oligonucleotides are listed in Table S2. Because CBS10367 derived haploids tend to diploidize, all gene editing experiments were performed in a diploid background. This required two successive gene targeting experiments with different selective markers to modify the two alleles of a gene.

Growth conditions and media, as well as detection of meiotic DSBs by PFGE followed by Southern blot are as in (50). Briefly, YPD medium (yeast extract 1%, peptone 2%, glucose 2%) was used for vegetative growth at 30 °C and sporulation was induced on 1% potassium acetate. Inactivation of both alleles of the chitin synthase III gene *CHS3* is a prerequisite for tetrad dissection in *L. kluyveri*.

### CBS10367 genome sequencing

Genomic DNA from CBS10367 was prepared using Qiagen tips according to the manufacturer instructions and sequenced at BGI. Reads were mapped to the *L. kluyveri* reference genome (downloaded from http://gryc.inra.fr/storedData/download/Lachancea_kluyveri/CBS_3082/) with BWA (v.0.7.4), using mem option (89). Duplicated reads were marked with Picard-tools (v.2.10.5) (broadinstitute.github.io/picard) and local realignment around indels and variant calling were performed with GATK (v.3.7-0) (90). Default parameters were applied except for the realignment step (GATKIndelRealigner), in which the following parameters were set: ‘-maxReadsForConsensuses 500 -maxReadsForRealignment 40000 -maxConsensuses 60 -maxPositionalMoveAllowed 400 -entropyThreshold 0.2’. Variants were called with GATK HaplotypeCaller function, before running GATK Variant Annotator to add allele balance information in the vcf file.

The complete genome sequence of the CBS10367 isolate was inferred both from the homozygous variants (SNPs and indels) and specifically from the homozygous SNPs (by removing indels with --remove-indels option from vcftools) into the reference sequence with GATK FastaAlternateReferenceMaker. This yielded the CBS10367_SNP_inferred and CBS10367_SNP_indels_inferred genome sequences. For SNPs and indels density analysis along chromosomes, the following variants were excluded: those located in the subtelomeric regions (30 kb from the beginning/end of the chromosome), and heterozygous variants for which the alternative residue *vs* (reference + alternative residues) allele ratio deviates from 0.4 to 0.6.

### CC-seq experiments

The *sae2* CBS10367 strain (LAKL 221) was used to perform meiotic time courses. 3 x 10^10^ cells were harvested for each CC-seq experiment six hours after meiosis induction, which corresponds to the peak of Spo11-DSBs (50). This amount of cells corresponds to ca. 10 times the amount of cells used for *S. cerevisiae* by (56) because of lower recovery yields in *L. kluyveri*. Likely because of different cell wall compositions, spheroplasting using zymolyase treatment takes longer in *L. kluyveri* compared to *S. cerevisiae*. We therefore used mechanical cell lysis instead of enzymatic lysis. Cells were lyzed in 2 ml screw cap tubes with 300 microliters of glass beads (425-600 micrometer of diameter, Sigma) per tube using a Fast prep (MP Biomedicals). We used three cycles of 30 seconds at 4 m/s, with five minutes on ice between the cycles. Cell lysates were harvested by centrifuging the screw cap tubes after drilling a hole at their base for 2 minutes at 950g at 4°C. From this point, cell lysates were treated as in (56). The efficiency of the CC-seq purification procedure was confirmed by Southern blot analysis of the previously characterized GPI18 Spo11-DSB hotspot using PflMI digested DNA (Figure S1A).

### CC-seq data analysis

Sequences were analyzed as in (56). *L. kluyveri* reference genome is CBS10367_SNP_inferred. The terminalMapper script from (56) generates a file with four columns corresponding to (i) the chromosome, (ii) the coordinate, (iii) the number of DSBs mapped on the Watson strand at this position and (iv) the number of DSBs mapped on the Crick strand at this position. The number of DSBs mapped on the Watson and Crick strands showed a good correlation and were pooled together for our subsequent analyses.

Unlike *S. cerevisiae* CC-seq data, *L. kluyveri* CC-seq data contained important background signal (about 60% of the entire signal) that needed to be filtered to extract the specific signal. We empirically determined a background signal threshold after signal smoothing with a Hann sliding window function over 15 units, a unit being a coordinate with a number of associated DSBs >0. Since intragenic regions are expected to be depleted in specific signal, we considered the mean signal per base pair over the entire intragenic regions as the background threshold that we subtracted to any genomic position. Only positive signals were kept for subsequent analyses. This denoising procedure suppressed around 81% and 73,5% of the total smoothed signal in experiment #1 and 2, respectively (Figure S1B). Experiment #2 showed the best enrichment. The number of DSB counts per bp is normalized by the total number of DSB counts per experiment and expressed as Hit per Million Hits mapped (HpM). All the analyses were performed using R (3.5 version or later) and RStudio (1.1.456 or later). Spo11-DSBs hotspots detection was performed using SAM files with MACS2 (2.1.1.20160309 version) (57) and the following parameters: callpeak -f SAM -g 11.345814e6 --nomodel --extsize 100 --shift 50 -q 0.01 for *L. kluyveri* and -g 12.157105e6 for *S. cerevisiae*.

The randomizations were performed by shuffling the Spo11-DSB counts associated to the intergenic regions in each dataset 1,000 times using the sample command from R. 1,000 comparisons between the randomized Spo11-DSB counts were performed, and the corresponding “Pearson” correlation coefficient r was calculated.

### Onholog analysis

Spo11-DSB counts of S288C *S. cerevisiae* strain were extracted from (10) supp data “FullMap.Spo11.S288C_chrX.txt”. The sum of the DSBs within intergenic regions was determined using the S288C reference genome (https://www.yeastgenome.org, downloaded March 2023). Pairs of syntenic intergenic regions flanked by ohnologs were identified with the list available at http://wolfe.gen.tcd.ie/ygob (60).

### ChIP-seq experiments

ChIP-seq experiments were performed as in (47, 91). All immuno-precipitations used the mouse anti-FLAG M2 antibody (Sigma). The expression of the FLAG-tagged proteins was verified by western blotting (Figure S4). ChIP-sequencing was performed with a Hiseq2500 sequencer (Illumina) using a 50-nt paired-end protocol. Two independent replicates of each condition were analyzed except for Zip3-FLAG that contained five replicates (Figure S6).

### ChIP-seq data analysis

#### Alignment

Sequencing quality control was determined using FastQC tool (http://www.bioinformatics.babraham.ac.uk/projects/fastqc/) and aggregated across samples using MultiQC (v1.7) (92). Reads with Phred quality score less than 30 were filtered out. Paired-end reads were mapped to CBS10367_SNP_inferred reference genome using Bowtie2 (v2.3.4.3) (93) with default parameters. Only mapped reads were filtered in using samtools (v1.9) (94) (*view*; *--exclude-flags 4*). Duplicate tags were removed using Picard Tools (v2.20.2) (https://broadinstitute.github.io/picard/) (*MarkDuplicates*; *REMOVE_DUPLICATES=true*) and mapped tags were processed for further analysis. Reads mapping to the mitochondrial genome and ribosomal DNA were removed from any analysis.

#### BigWig files

For visualization purpose, mapped tags were converted into bigwig and normalized by their library size using the deepTools (v3.2.1) (95) (*bamCoverage – scaleFactor library_size_factor –binSize 5*). Reads were extended using values previously computed by deepTools (*bamPEFragmentSize*) and used through the *– extendReads* option. Untagged-subtracted bigwig files were computed from the normalized bigwig files using deepTools (*bigwigCompare –operation subtract*). BigWig files from multiple replicates were merged into a single file using a custom bash script (*bigWigMerge, bedGraphToBigWig*). All sequencing data were visualized using the Integrative Genomics Viewer (IGV; v2.5.2) (96). Immunoprecipitated over untagged ratio profiles were generated with bamCompare using the –operation ratio (Figure 3B).

#### Peak calling

High confidence binding sites were determined using MACS2 callpeak (v2.1.2) (57) in broad mode (--broad; --broad-cutoff 0.05; --qvalue 0.01). Each sample’s fragment size was previously computed by deepTools (bamPEFragmentSize) and used through the --extsize option. Untagged strains at the same timepoint were used as controls.

#### ChIP-seq pileup analyses

ChIP-seq signals were piled up around specific features by processing bigwig and untagged-substracted bigwig files through the deepTools (*computeMatrix; --referencePoint center; --binSize=50*) and using custom R scripts for visualization. Background signals from plots containing signals of experiments done by immunoprecipitating different proteins have been adjusted for better comparison (Fig S7C). The background signal of a given condition was determined as the average of the extreme left and right values of the plot profile. The differences between background signals were used to correct signals by making their background levels similar.

#### Venn diagrams

Venn diagrams analyses were performed from peak coordinates bed files obtained from MACS2 peak caller and using python matplotlib_venn package (v0.11.5) (https://github.com/konstantint/matplotlib-venn) for visualization. Peaks were intersected using bedtools *intersect* (v2.28.0) (97) with a minimum overlap of 1-base. Overlaps significance was determined by hypergeometric test (*phyper*; *lower.tail = false*). Since a given peak can overlap multiple peaks of other proteins, some peaks were merged during the intersect processes leading to small variations in total peak numbers.

### Hi-C

Hi-C experiments were performed using the CBS 10367 isolate. To harvest enough cells per time point (ie ca 2 x 10^9^ cells), four independent 200 ml cultures in 2 l flasks were used. Equal samples were harvested from the four flasks at 0, 2, 3, 4, 5, 6, 7 and 8 hours post meiosis induction and treated with 3% formaldehyde for 20 minutes under constant shaking. Samples were quenched using 0.35M glycine at 4°C for 20 minutes. Cells were harvested by centrifugation at 4000 g for 4 minutes at 4°C, washed once with potassium acetate 1% (sporulation medium), transferred to 1.5 ml tubes and pellets were flash frozen using liquid nitrogen.

### Generation and normalization of contact maps and ratio maps

Alignment of paired-end reads and processing of the contact data were done with Hicstuff pipeline on the CBS10367_SNP_inferred reference genome, using with the following parameters: aligning with bowtie2, filtering out spurious 3C events and PCR duplicates based on read positions. Contact maps were normalized with the ICE function as described in (98) and generated with the ‘view’ mode of Hicstuff (99). Contact maps were binned at 10 kb for the whole genome or 1 kb for single chromosomes.

### Contact probability as a function of the genomic distance

Contact probability as a function of genomic distance P(s) and its derivative were computed using the ‘distance law’ function of Hicstuff on the following ∼1Mb chromosomal arms : Lakl0A-L, Lakl0B-R, Lakl0C-L, Lakl0E-R and Lakl0G-L arms.

### Generation of aggregated conhesin-centered contact maps and spectrum of loop score as a function of distance

Loops at the intersection of cohesin enrichment sites located on ∼1Mb of Lakl0A-L, Lakl0B-R, Lakl0C-L, Lakl0E-R and Lakl0G-L arms were scored with chromosight in quantify mode. Lowess-smoothed loop scores were plotted as a function of loop size data.

Cohesin pile-up contact maps were generated with chromosight (67) dividing averaged 29 kb windows centered on pairs of cohesin-enriched sites by the average of 10 windows centered on 10 random pairs of positions maintaining the same inter-distance distribution. Cohesin peaks whose enrichment determined by MACS2 was higher than 2 were selected.

### Meiotic chromosome spreads

Chromosome spreads from meiotic cultures were prepared from ∼6×10^8^ cells harvested at the indicated time-points following the protocol described in (100) with some modifications. The addition of 0. 005% of Antifoam (Sigma) to the pre-sporulation and sporulation media helped avoid the clustering of cells in the cultures. Yeast cells were first processed into spheroplasts by incubation at 30°C for 15 (for *S. cerevisiae* strains) to 40 (for *L. kluyveri* strains) minutes with (400 µg/ml final) Zymolyase 100T (120493-1, AMS Biotechnologie). Spheroplasts were washed and resuspended in cold MES(2-(N-morpholino)ethanesulfonic acid)/sorbitol buffer to which equal volumes of 100mM sucrose solution was added and incubated for at least 10-30 minutes before spreading.

For chromosome spreading, slides were divided into two squares using an ImmEdge hydrophobic pen (Vector Labs) and placed horizontally in a humid chamber followed by the addition of 10 μl of cell suspension per square. The cell suspension was subsequently lysed with 1% NP40 (IGEPAL CA-630, Sigma) and fixed with 3% paraformaldehyde (PFA) containing 3.4% sucrose and 0.04% Sodium Dodecyl Sulfate (SDS). In addition, 0.001M dithiothreitol (DTT) was added to MES/sorbitol buffer, sucrose and PFA solutions. The slides were gently swirled for homogenization, before incubation in the closed humid chamber overnight. The humid chamber was opened for a few hours the next day to allow almost complete drying of the cell suspension.

### Immunostaining

Slides with spread nuclei were sequentially incubated with the proper primary and secondary antibodies. Slides were washed before and in the intermittent steps in a Coplin jar for 10 min each in PBS-Tween 0.05% containing 0.24% Photo-Flo (Kodak) and 0.01% Triton X-100, in that order, and 10 min in 0.01% Photo-Flo aqueous solution on a shaker. Primary antibodies used were affinity purified rabbit anti-*S. cerevisiae* Zip1 antibody (Gift from A. Macqueen, dilution 1:100), mouse monoclonal anti-FLAG primary antibody M2 (Sigma, dilution 1:300). Secondary antibodies used were Alexa Fluor488 donkey anti-mouse (Life technologies, A21202), Alexa Fluor488 goat anti-rabbit (A-11008, Thermo Fischer Scientific), Alexa Fluor568 goat ant-imouse (A-11004, Thermo Fischer Scientific), Alexa Fluor568 donkey anti-rabbit (Life technologies, A10042), Alexa Fluor647 donkey anti-rabbit (Abcam, ab150067), Alexa Fluor647 donkey anti-mouse (Abcam, ab150107), all at 1:200 dilutions. Slides were mounted using an anti-fade mounting medium with DAPI (Vectashield with DAPI, H-2000-10, Vector Laboratories).

### ImmunoFISH and DNA probe preparation

Immunostained chromosome spreads were re-fixed using PFA 2% in PBS for 10 min then washed twice in PBS for 5 min. Slides were then dehydrated for 4 min in ethanol 70%, 4 min in ethanol 90% and 5 min in ethanol 100% before air drying for 10 min. To generate FISH probes, we used the nick translation kit from Abbott Molecular following the manufacturer’s instructions and using CF dye-conjugated dUTP (Biotium), on a pool of PCR products. We pooled 4 PCR products for the Lakl0C-left and 3 PCR products for the Lakl0C-Right, each being about 6 kb long. For the rDNA region we used 1 PCR product of 8 kb. Each product was amplified using LongAmp Taq polymerase (NEB #M0323L). To avoid non-specific hybridization of the Lakl0C-left and Lakl0C-right probes to the rDNA locus, slides were digested with RNase A (Sigma Aldrich/MERCK) during the secondary antibody incubation, and rDNA probes (labelled or not) were always hybridized during FISH.

Labeled DNA (500 ng) was precipitated during 30 min incubation at −20°C after adding 0.5 volume of 7.5 M ammonium acetate and 2.5 volumes of cold 100% ethanol. After washing with 70% ethanol and air drying in the dark, the pellet was dissolved in 15 μl of hybridization buffer. Chromosomal DNA and probes were co-denatured for 6 min at 78°C and left overnight at 37°C. Washes were performed using PBS-Tween 0.1% for 5 min twice and once in SSC 0.4X-NP40 0.3%. Finally, slides were air dried and mounted using Vectashield with DAPI.

### Image Acquisition

Fluorescence microscopy was performed on a ZEISS IMAGER Z2 upright microscope, equipped with a CoolSNAP HQ² camera (PHOTOMETRICS, roper Scientific), DAPI, GFP, TEXAS red and Cy5 filter sets and controlled by METAMORPH Software. Images were acquired using a 100X/1.46 NA oil immersion objective and a tubelens at 1.25X. Images shown are projections of two to three individual focal planes along the z axis.

### Measurement of chromosome axis length and DAPI width

Original, unmodified spread images of pachytene nuclei (DAPI-stained regions) were used to measure axis length and DAPI width. The measurements were done manually as described in (84). They showed that the chromosome axes can be reliably visualized and measured by immunostaining either Zip1 or Rec8 as their abundance are highly correlated in the synapsed regions during pachytene. In this study, we used Zip1 length as a proxy for axis length measurement and DAPI width as a proxy for the loop-size estimation.

Image analysis and quantification was performed using the Fiji software. Zip1 length was measured by tracing its immunofluorescence signal. The beginning and the end of each Zip1 line is determined manually from the pixel intensity at the edges of each line. For DAPI width estimation, a straight line perpendicular to the DAPI-stained chromosome fragment, covering its entire width, was drawn at three different locations (middle and closer to the ends) on the same chromosome. The width of each chromosome was calculated as the average length of three perpendicular lines drawn across it. DAPI signal intensities of the pixels along each perpendicular line were used to differentiate the background from the exact length of the line spanning the width of the chromosome. Graphs and statistical analyses were performed using GraphPad Prism 9 software.

### Accession numbers

The Illumina reads related to the CBS10367 whole-genome sequencing are available in the Sequence Read Archive under the BioProject PRJEB60460.

The CC-seq, ChIP-seq and Hi-C data discussed in this publication have been deposited in NCBI’s Gene Expression Omnibus (101) and are accessible through GEO SuperSeries accession number GSE237708 (https://www.ncbi.nlm.nih.gov/geo/query/acc.cgi?acc=GSE237708).

To review GEO accession GSE237708: Go to https://www.ncbi.nlm.nih.gov/geo/query/acc.cgi?acc=GSE237708 Enter token mnkngwuctpepvsb into the box.

---

**Figure S1:**
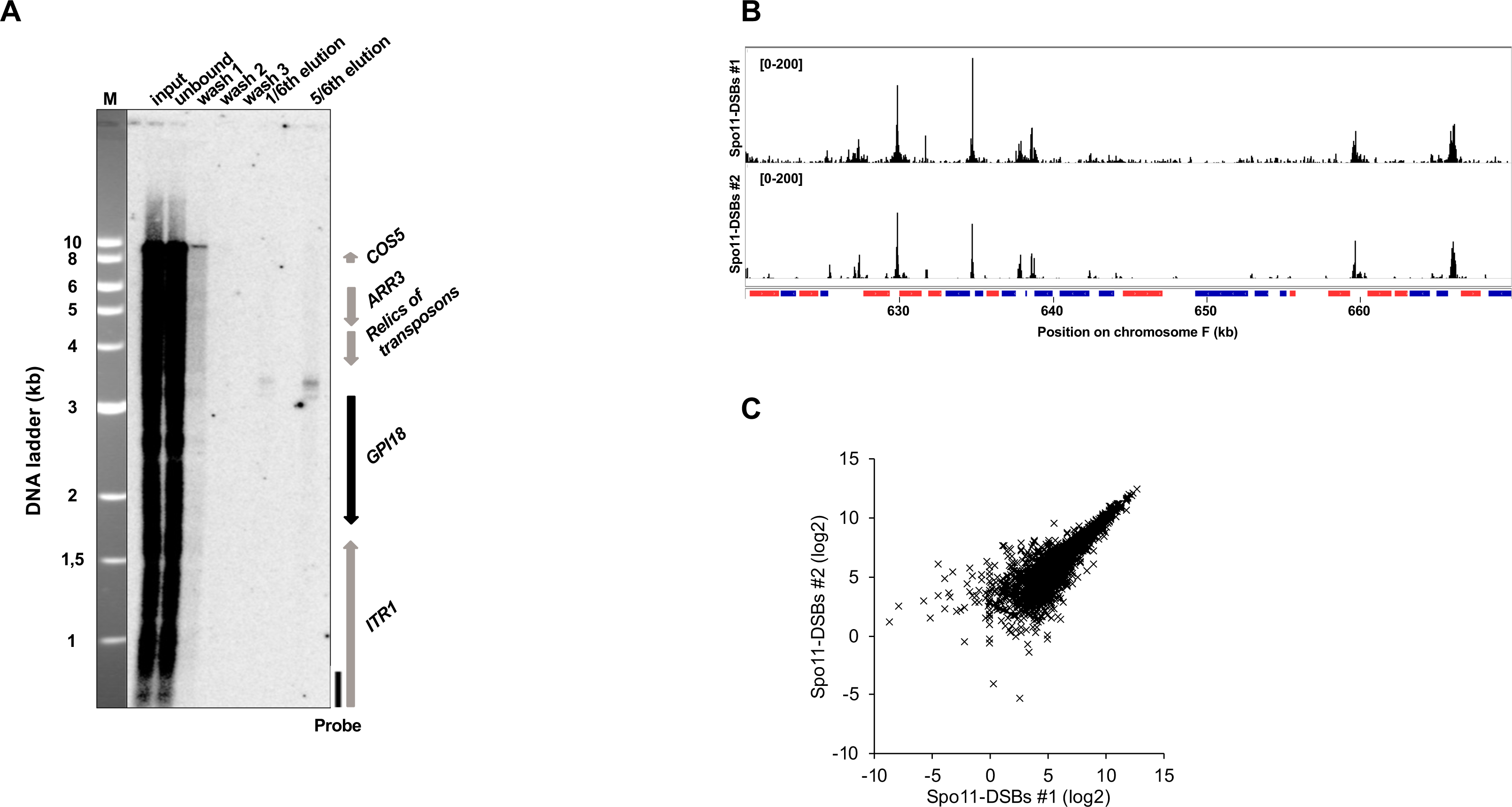
Spo11-DSB identification by CC-seq. **A.** Test for locus specific enrichment of Spo11-DSBs. The CC-seq Spo11-DSB purification protocol was applied to 6.5 x 10^8^ *L. kluyveri* cells at six hours post meiosis induction. Here, the DNA was digested by PflMI instead of sheared by sonication. Aliquots of the indicated fractions were analyzed by 0.8 % agarose gel electrophoresis followed by Southern blot using the indicated probe (vertical black bar) as in (50). The right side of the blot shows the gene (arrows) organization around the *GPI18* locus with arrow heads indicating the relative transcription orientations. M: molecular weight ladder (kb). A Spo11-DSB signal specific to the GPI18 promoter is detected in the elution fractions. **B.** Two independent CC-seq profiles after background removal as in Figure 1B. The signal/noise ratio of the top experiment is lower than that of the bottom experiment. Only the bottom experiment was kept for further analyses. **C.** Correlation of the two independent CC-seq experiments after background removal inside the intersected hotspots detected by MACS2 (n = 2038). Normalized in hit per million reads of the corresponding sequencing library.

**Figure S2:**
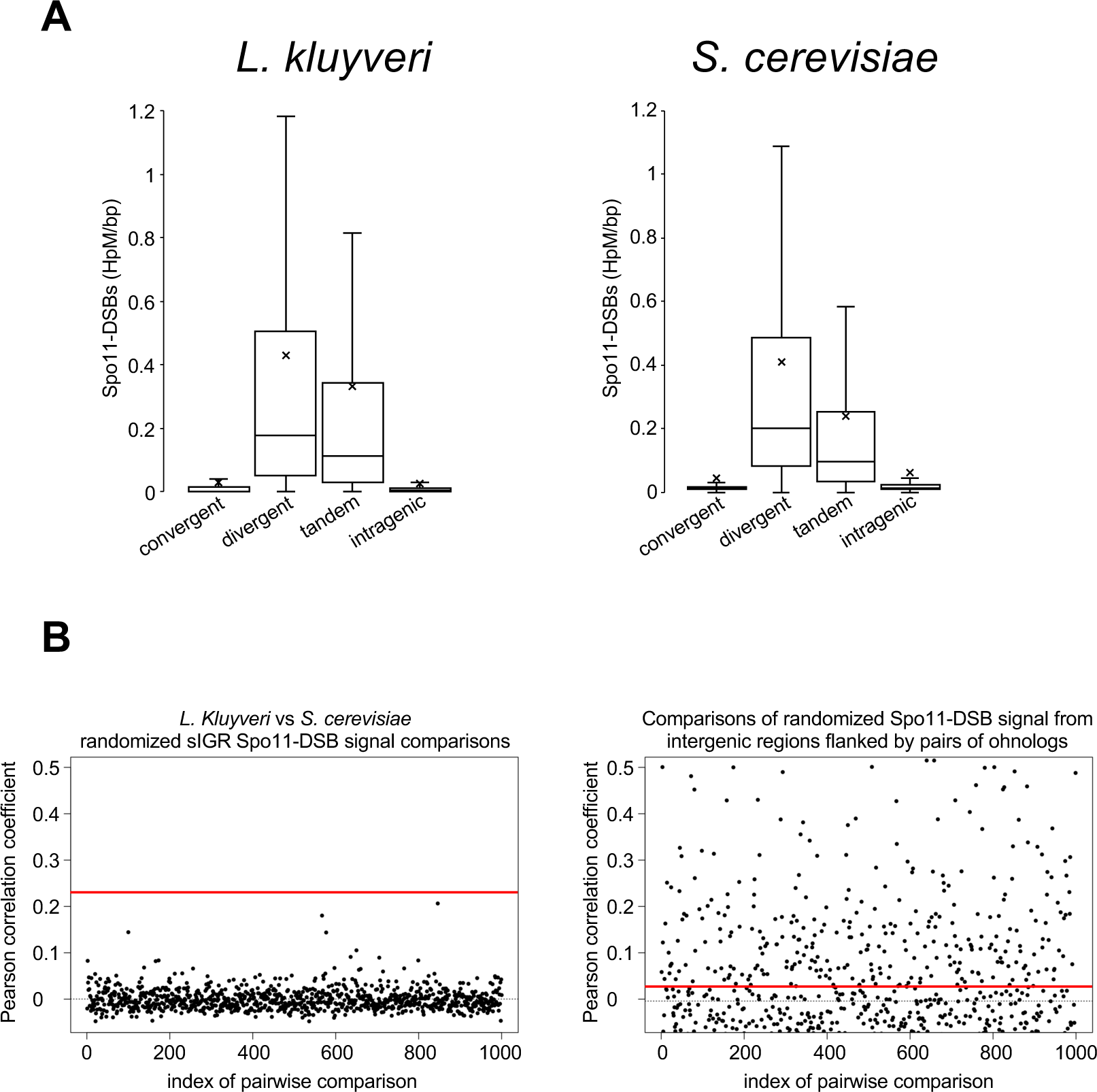
Characteristics of Spo11-DSB signal. **A.** Box plot representing the distribution of the CC-seq signal (mean DSB count in hits per million reads of the sequencing library per kb) according to the type of genomic location as in Figure 1B in *L. kluyveri* and *S. cerevisiae*. **B.** Pearson correlation coefficient of 1,000 pairwise comparisons of randomized Spo11-DSB signal from syntenic intergenic regions (IGR) between *L. kluyveri* and *S. cerevisiae* (left), and between *S. cerevisiae* intergenic regions flanked by pairs of ohnologs (right). The red lines represent the actual coefficients from Figure 2. The small-dots lines represent the mean coefficients from the comparisons of the randomized datasets.

**Figure S3:**
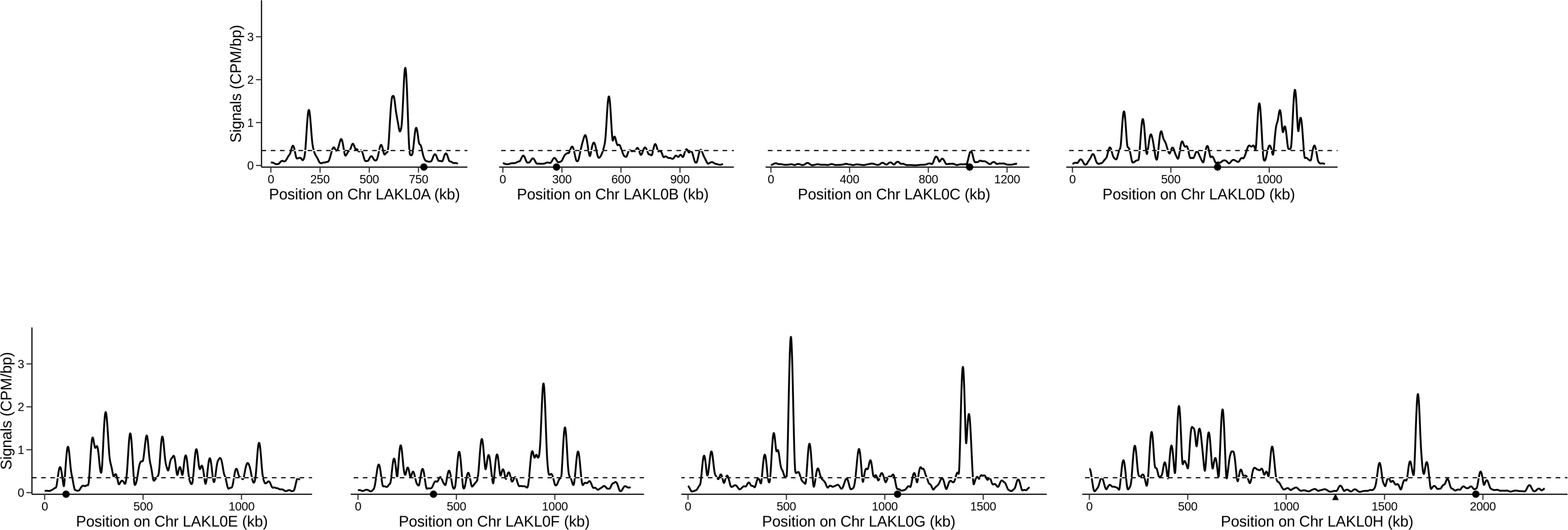
Spo11-DSB CC-seq chromosome-wide signal smoothed with a Hann sliding window of 50kb. The dotted lines represent the genome average.

**Figure S4:**
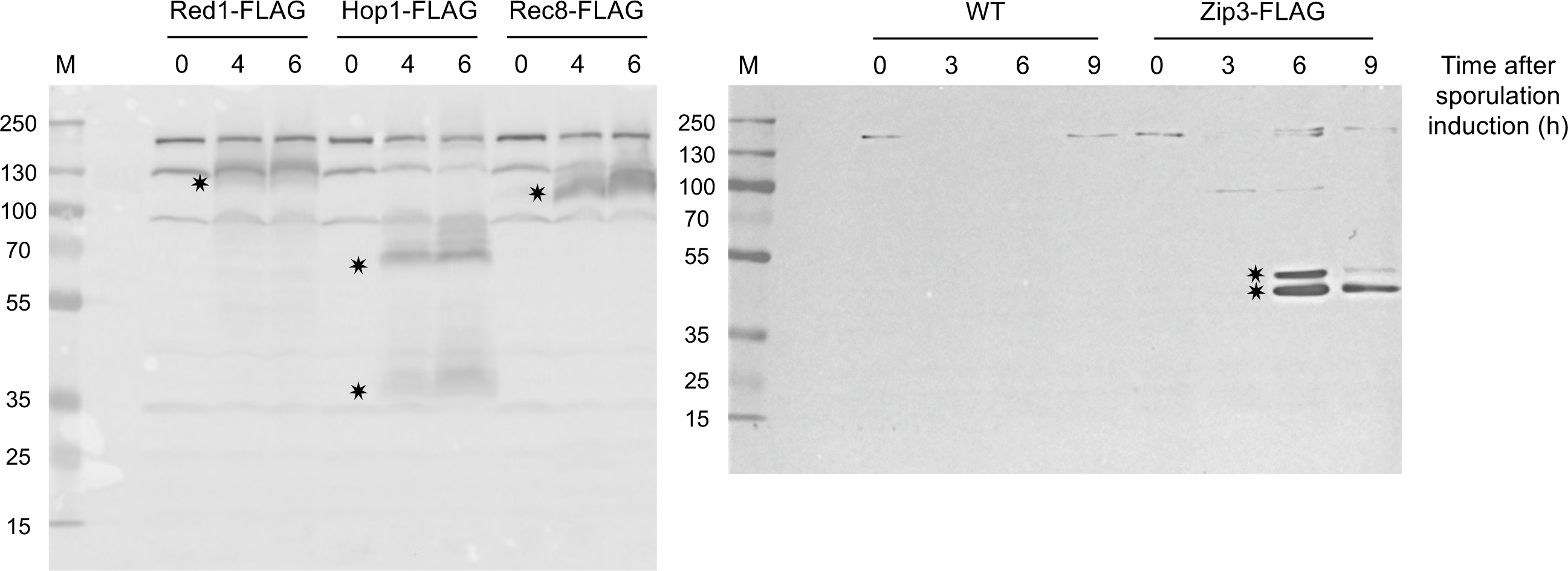
Western blot analysis of Red1, Hop1, Rec8 and Zip3. Protein extracts from cultures of the indicated time points after sporulation induction were run on acrylamide gels, transferred to a nitrocellulose membrane, incubated with an anti-FLAG antibody (mouse) which is common to the four proteins analyzed and revealed with an anti-mouse-HRP antibody. M: molecular weight marker with sizes in kDa indicated on the left. Estimated molecular weight are: 99 kDa for Red1, 69 kDa for Hop1, 79 kDa for Rec8 and 37 kDa for Zip3. Each of these proteins is fused with six histidines and three copies of the FLAG tag, which adds about 4 kDa.

**Figure S5:**
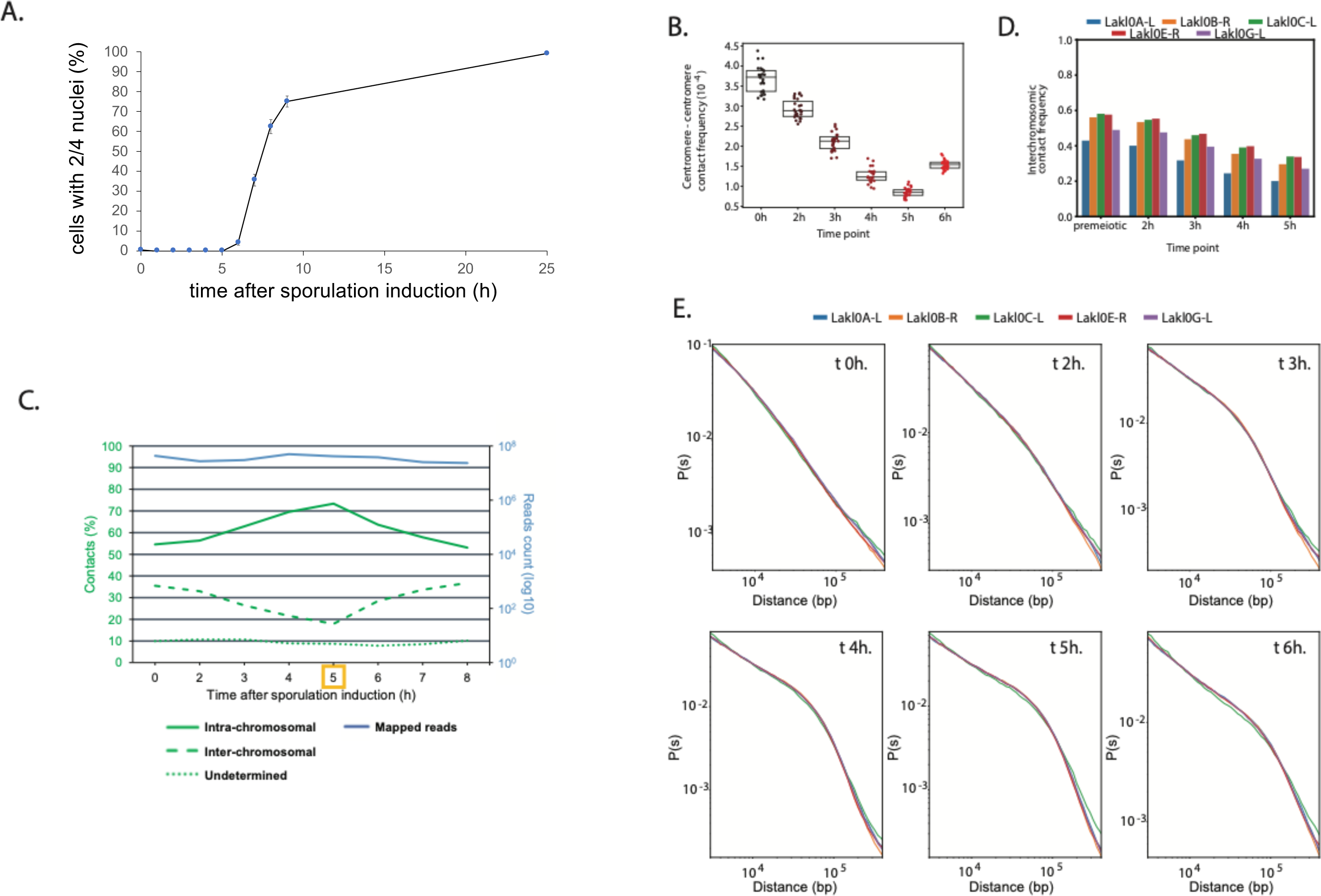
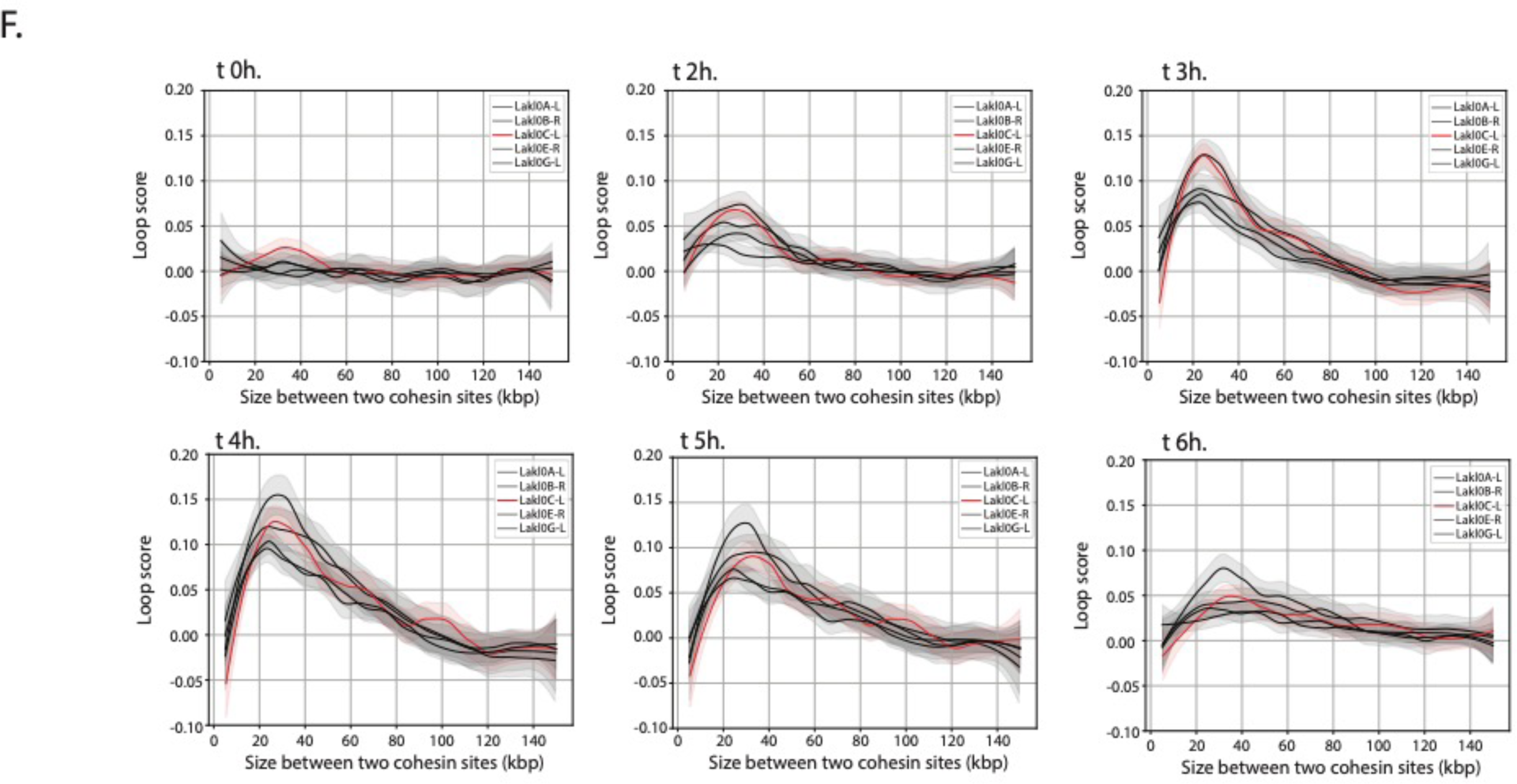
Meiotic progression and evolution of chromosomal contacts. **A.** Fraction of cells with two to four nuclei detected by DAPI staining as a function of time post meiosis induction (in hours). Error bars represent standard deviation among the four cultures pooled together for the Hi-C experiment. **B.** Inter-centromere contact frequencies at different time points of a meiotic time-course**. C.** The number of chromosomal contacts sequenced per each time point is measured on the right y axis. The left y axis represents the fraction of each type of contact, i. e. intra-chromosomal, inter-chromosomal or undetermined. **D.** Frequency of interchromosomal contacts for several ∼1Mb chromosome arms from zero to five hours of a meiotic time course. **E.** Distance laws of the several ∼1Mb chromosome arms from zero to six hours of a meiotic time course. **F.** Spectra of loop scores detected at the intersection of Rec8 peaks as a function of the distance separating the pairs of Rec8 peaks (kbp) for different time points of a meiotic time-course and for several ∼1Mb chromosomal arms.

**Figure S6.**
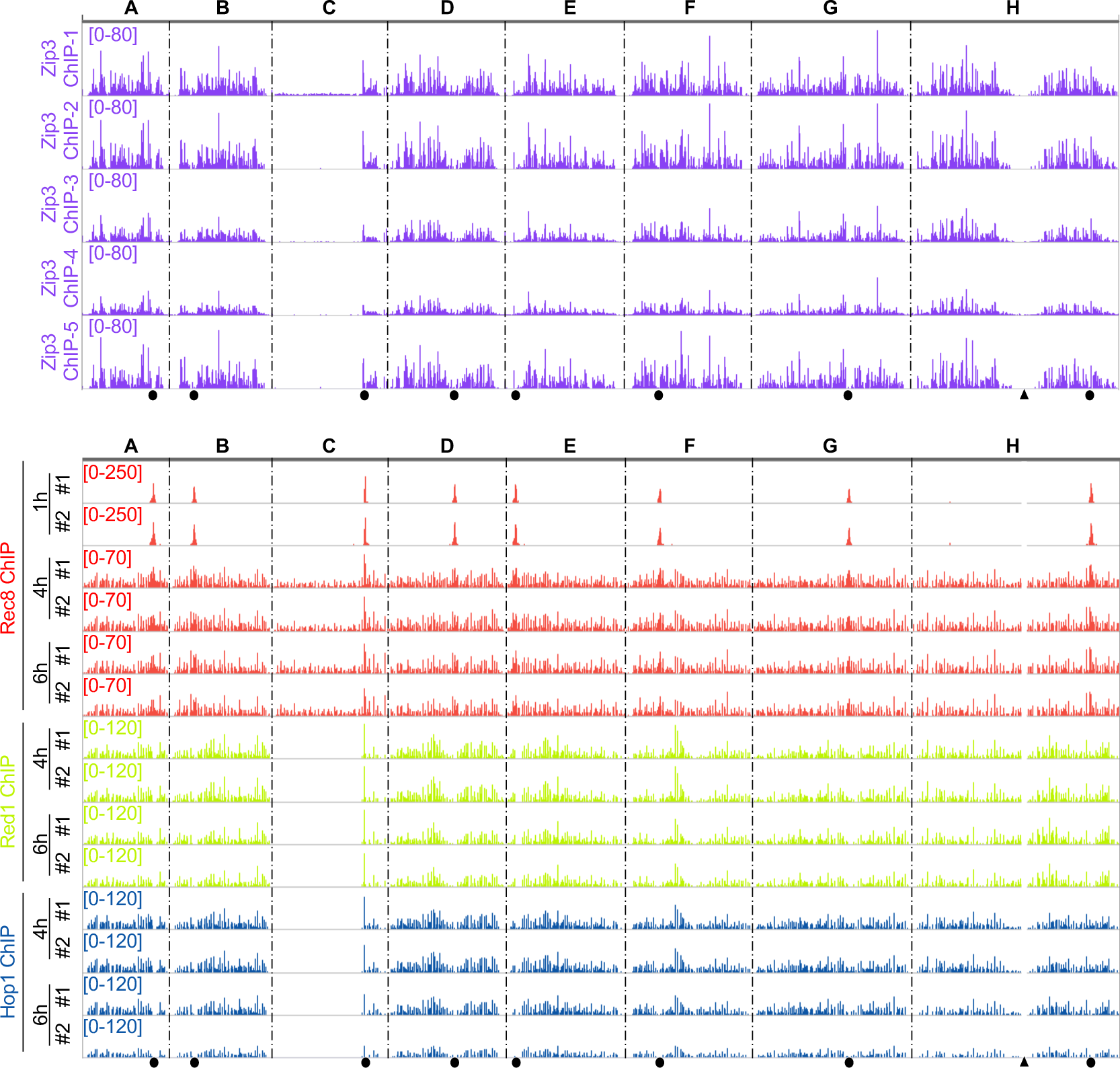
ChIP-seq individual replicates of Zip3-FLAG, Rec8-FLAG, Red1-FLAG and Hop1-FLAG. Same legend as in Figure 1A.

**Figure S7:**
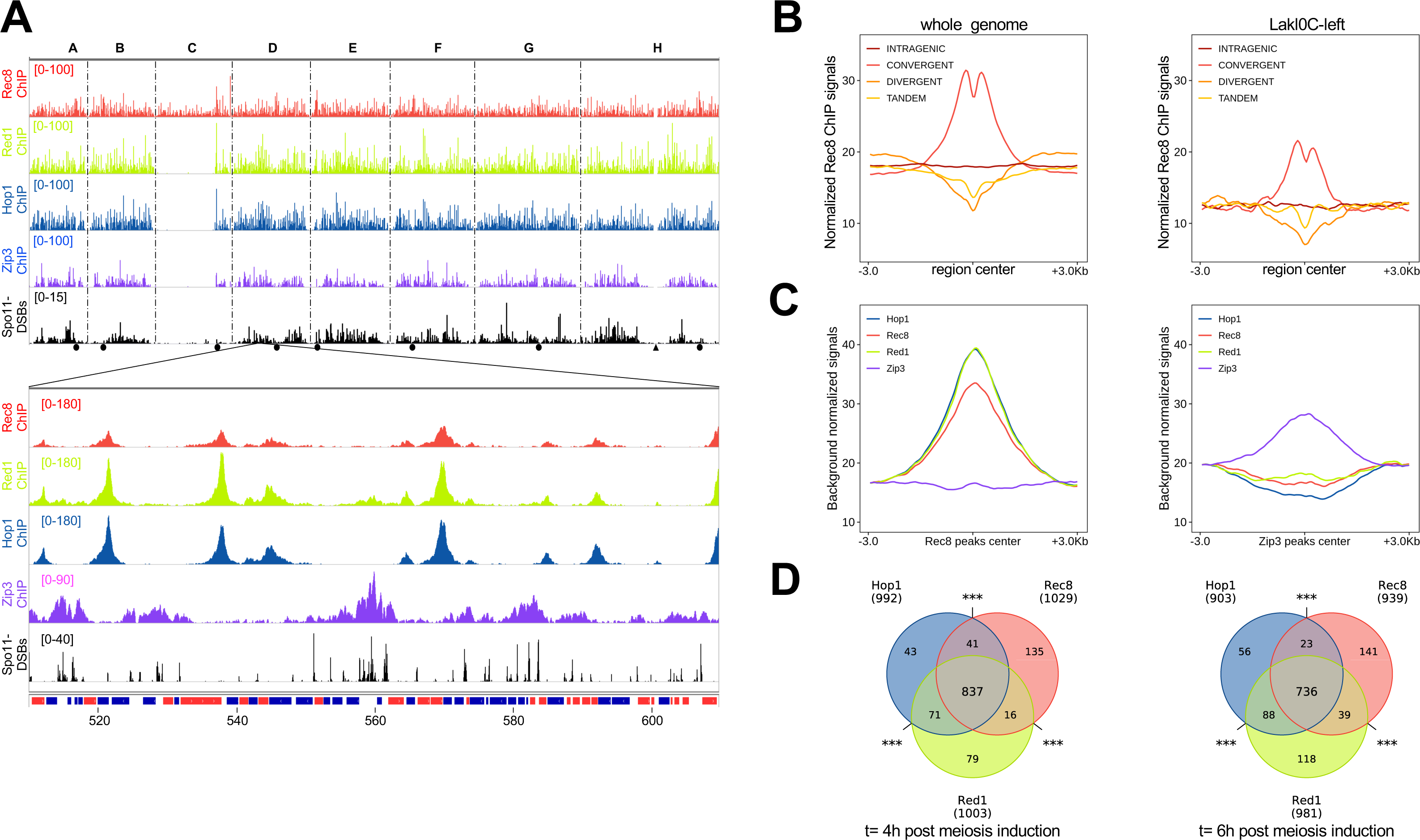
Comprehensive ChIP-seq analysis of the axis proteins Rec8, Red1 and Hop1, with the crossover marker Zip3 and the Spo11-DSBs. **A.** Combination of CC-seq data and ChIP-seq data from Figure 1A and Figure 4. ChIP-seq data are the means of all the replicates presented in Figure S6 (five replicates for Zip3 and two replicates for Rec8, Red1 and Hop1). **B**. Pileup of Rec8-FLAG ChIP-seq signals at six hours post meiosis induction overlapping intragenic regions, convergent intergenes, divergent intergenes, and intergenes in a tandem configuration, centered on the center of the region of interest. Left, whole genome; right, Lakl0C-left only. The lower Rec8-FLAG ChIP-seq signal on Lakl0C-left (right panel) likely results from a lower sequencing efficiency of this region due to an elevated GC content (see methods). **C**. Background-corrected pileup of Rec8-FLAG, Red1-FLAG, Hop1-FLAG and Zip3-FLAG ChIP-seq signals centered on Rec8-FLAG ChIP-seq peaks centers (left) and Zip3-FLAG ChIP-seq peaks centers (right). Rec8 and Zip3 peak coordinates overlapping rDNA coordinates where remove from the analysis. **D**. Overlaps between Hop1-FLAG, Red1-FLAG and Rec8-FLAG ChIP-seq peaks at four (left) and six (right) hours post meiosis induction. ***P < 0.001 (hypergeometric test).

**Figure S8:**
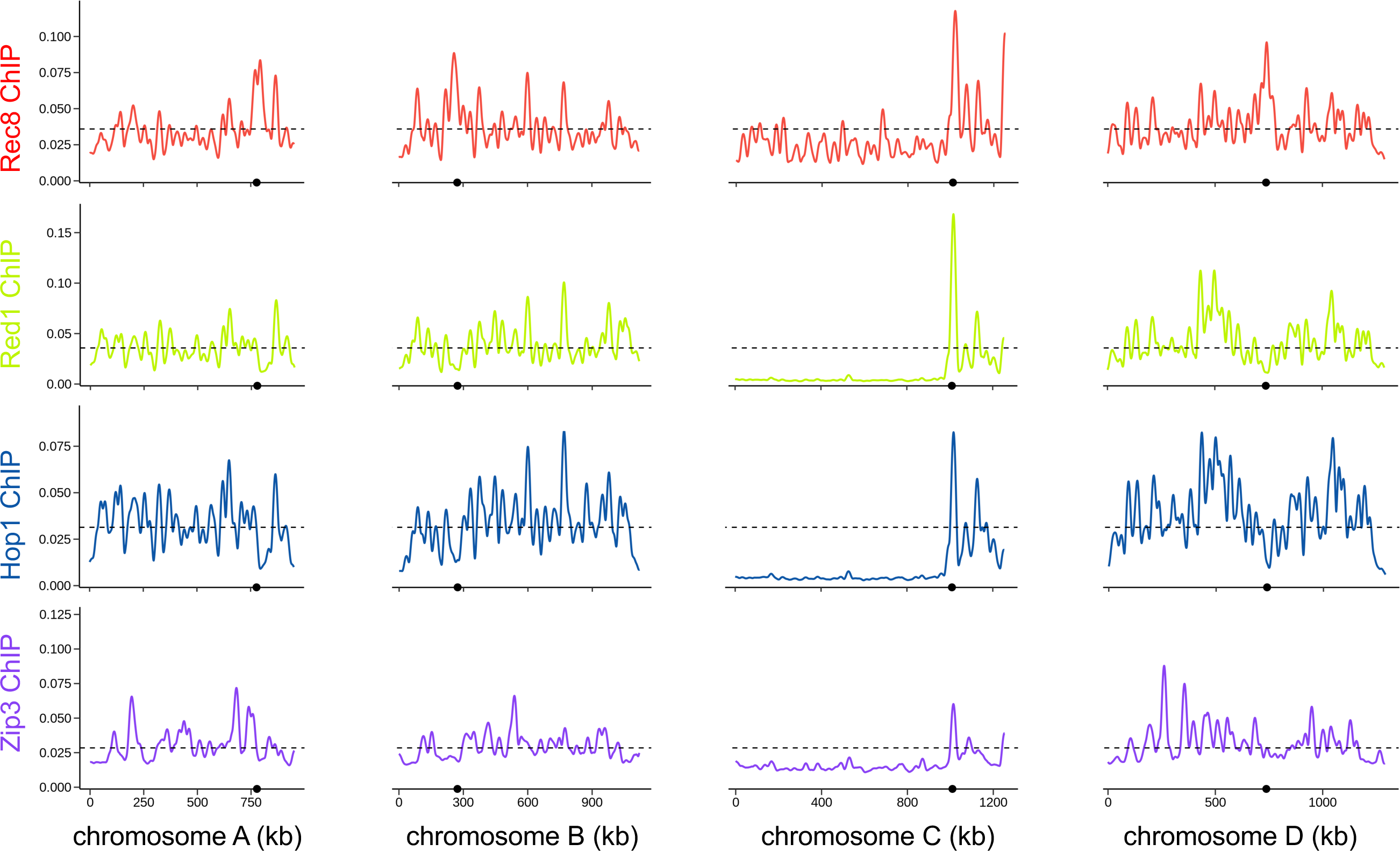
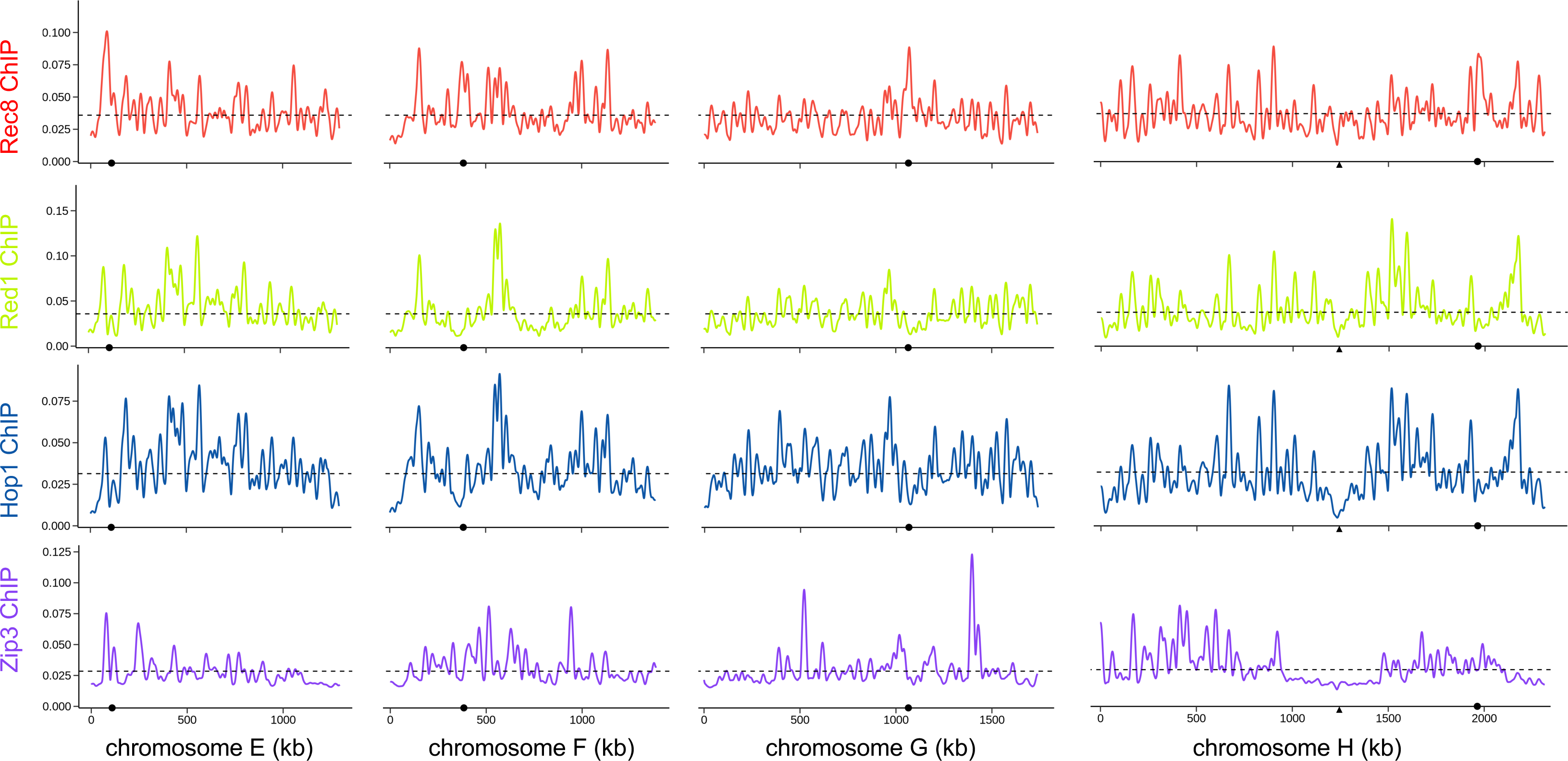
Rec8, Red1 and Hop1 ChIP-seq chromosome-wide signals smoothed with a Hann sliding window of 50kb. The dotted lines represent the genome average.

**Figure S9.**
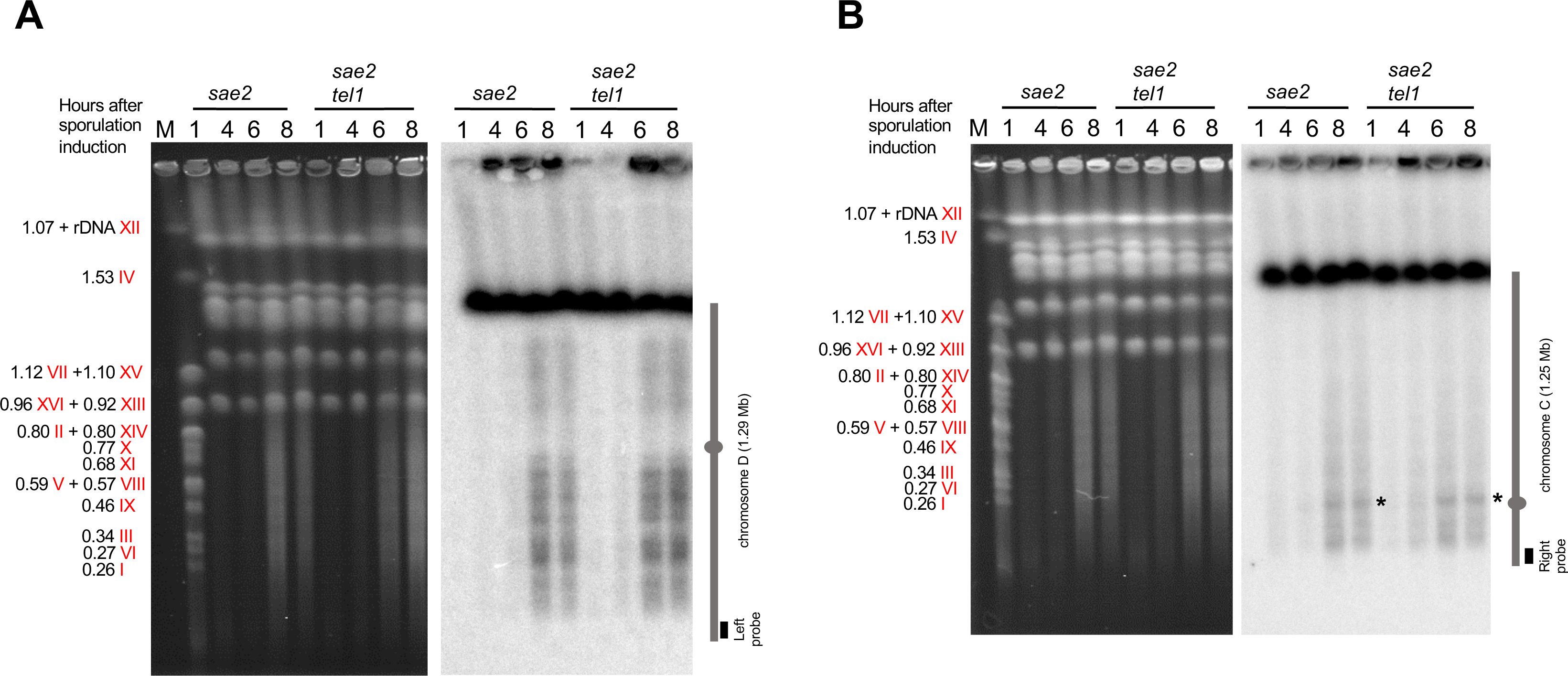
The absence of Tel1 does not restore Spo11-DSBs on Lakl0C-left. A. PFGE analysis of *L. kluyveri* meiotic chromosomes. Left: ethidium bromide-stained gel. M: *S. cerevisiae* SK1 chromosomes were used as molecular weight markers with their sizes indicated in Mb on the left. Right. Southern blot analysis using a chromosome D left probe. The vertical grey bar represents chromosome D and the grey oval indicates the centromere location. B. Same as in A except that a chromosome C right probe was used.

**Figure S10.**
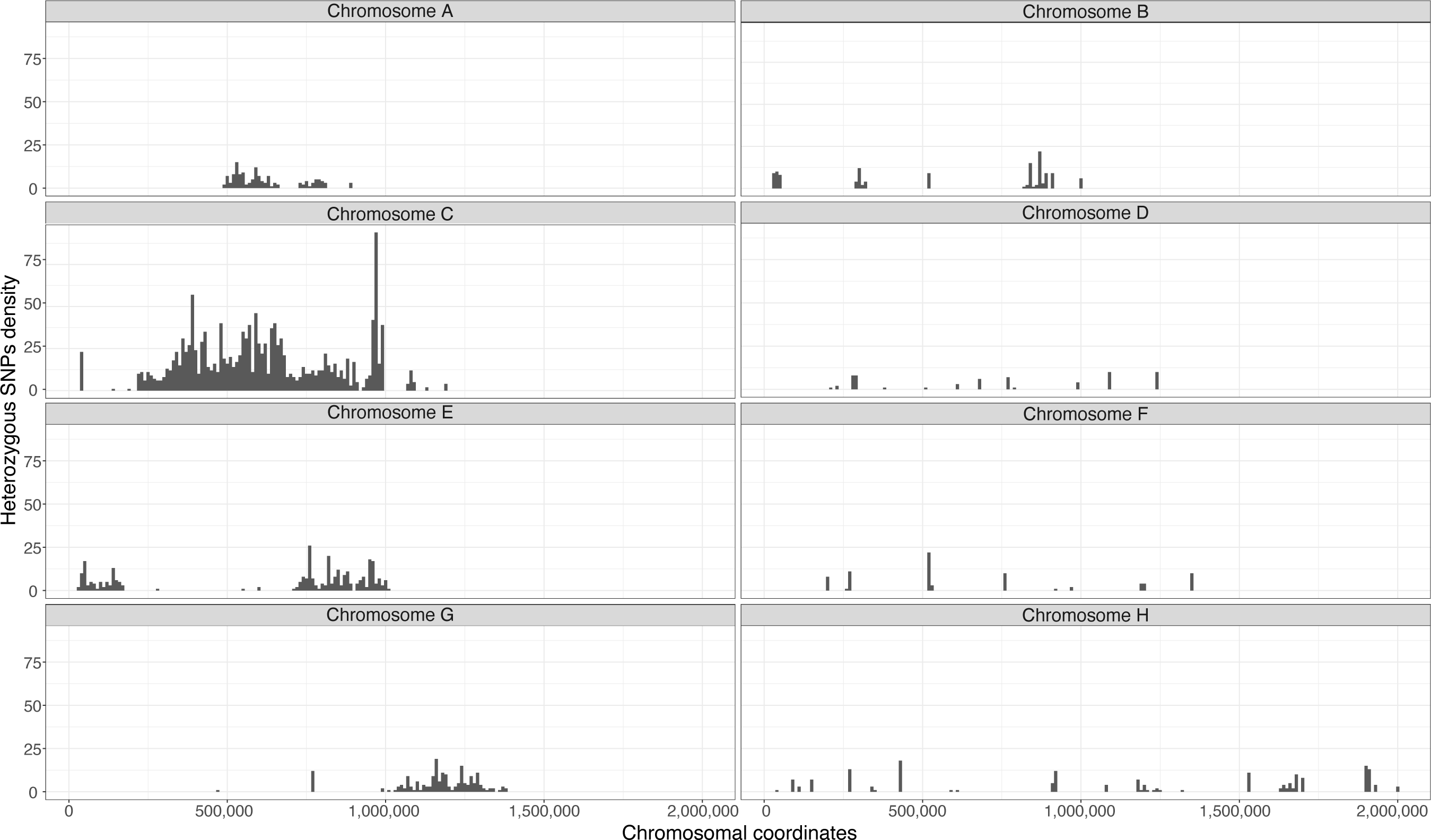
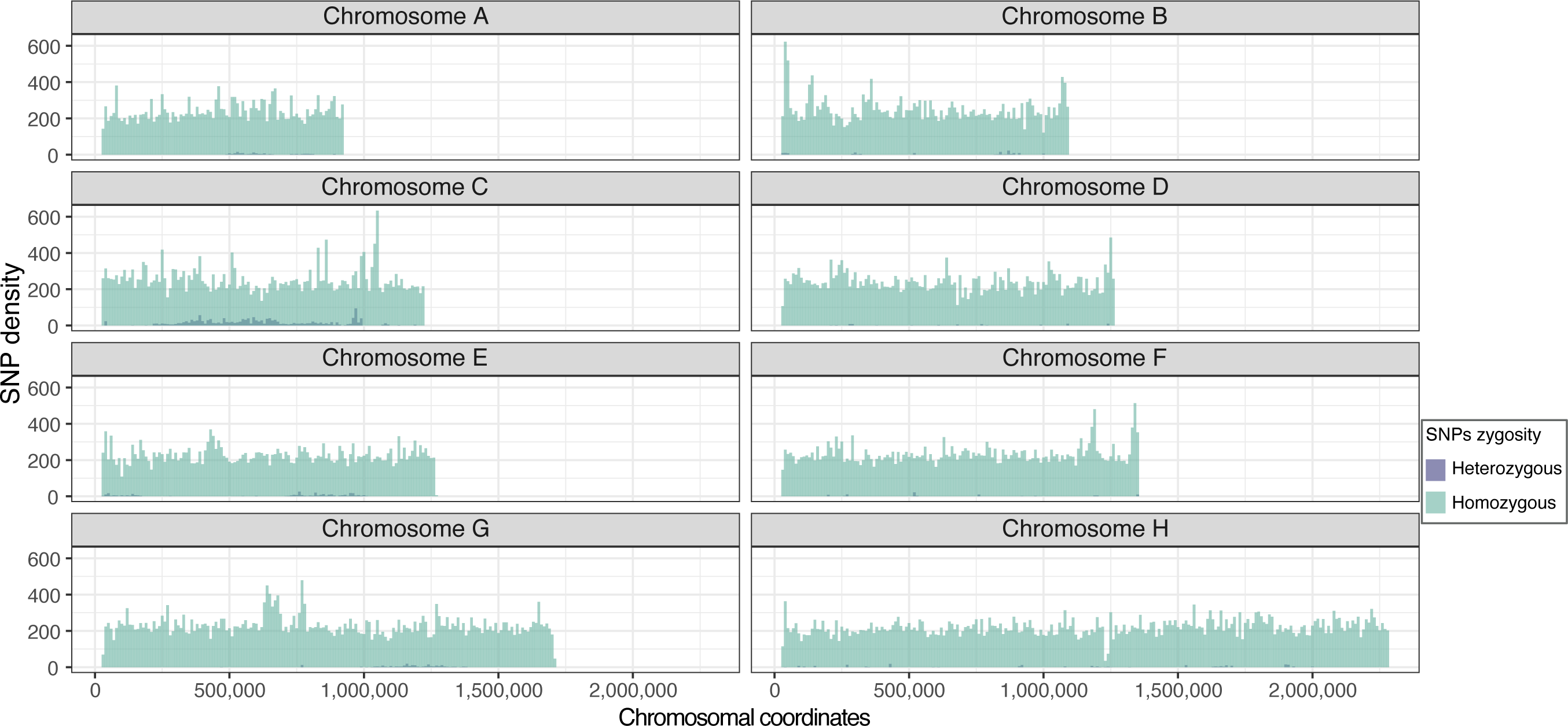
**A.** Number of heterozygous SNPs per 10 kb windows. **B**. Number of heterozygous (dark grey) and homozygous (light grey) SNPs per 10 kb windows.

**Table S1.**
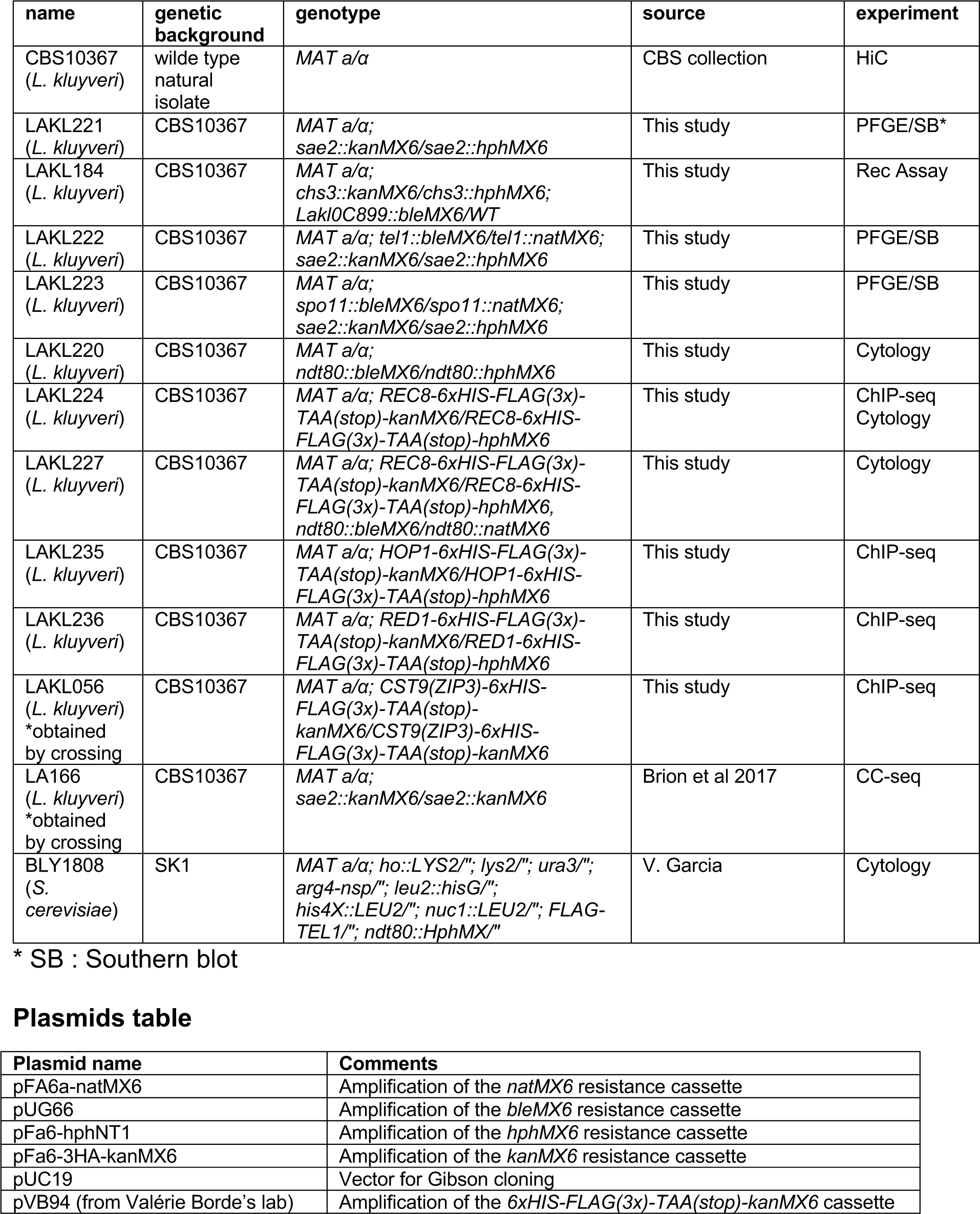
Yeast strains table.

**Table S2.**
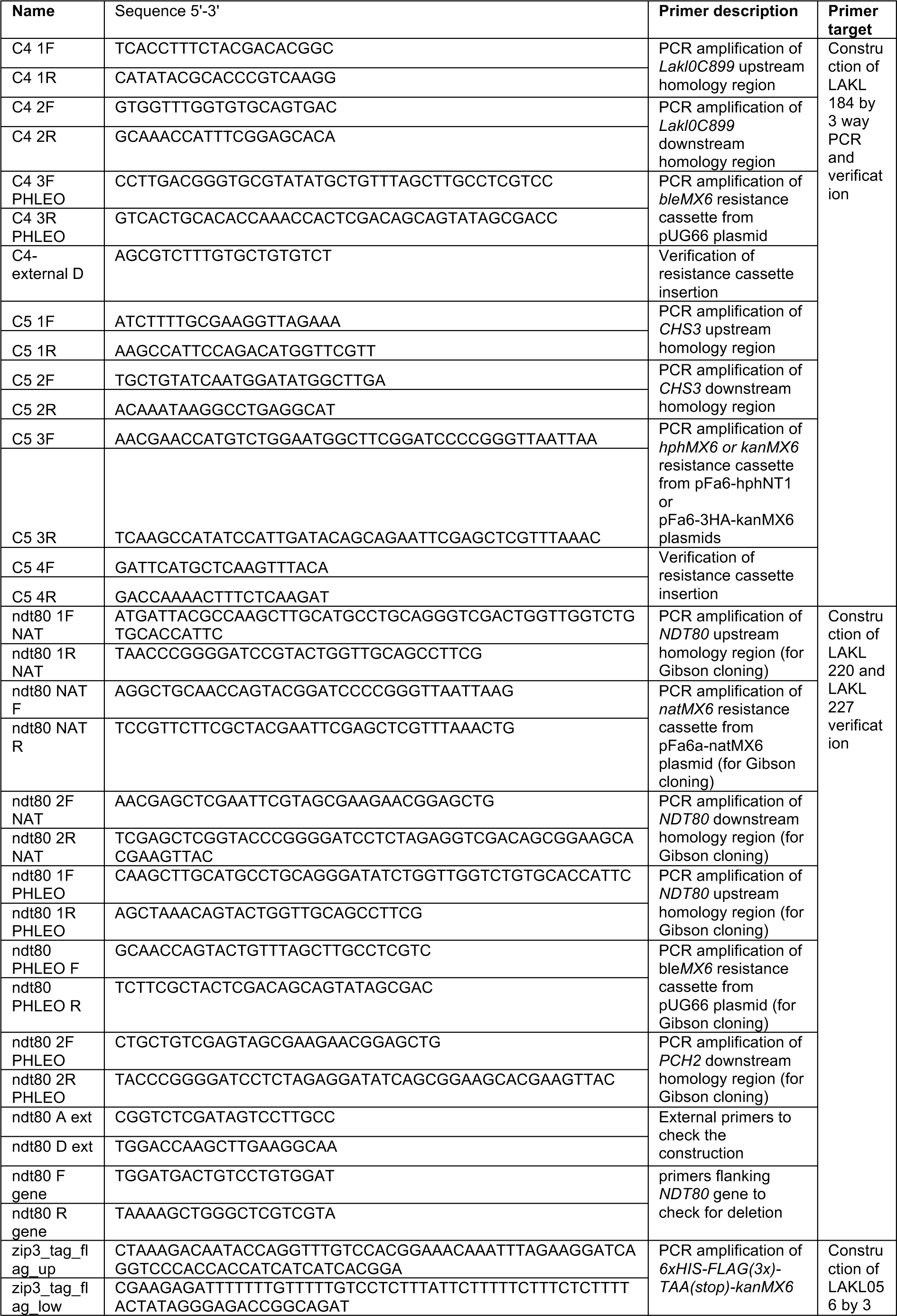
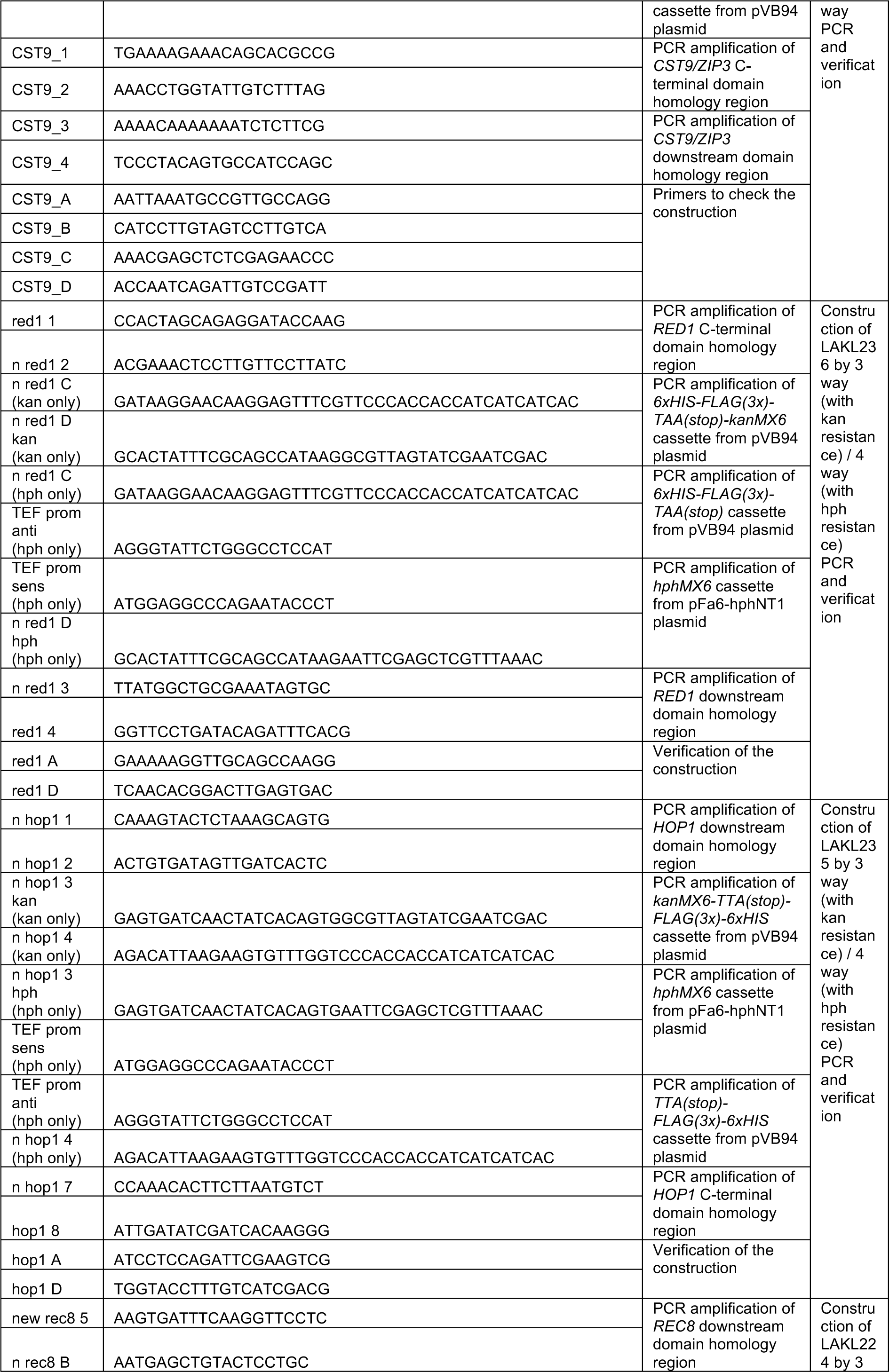
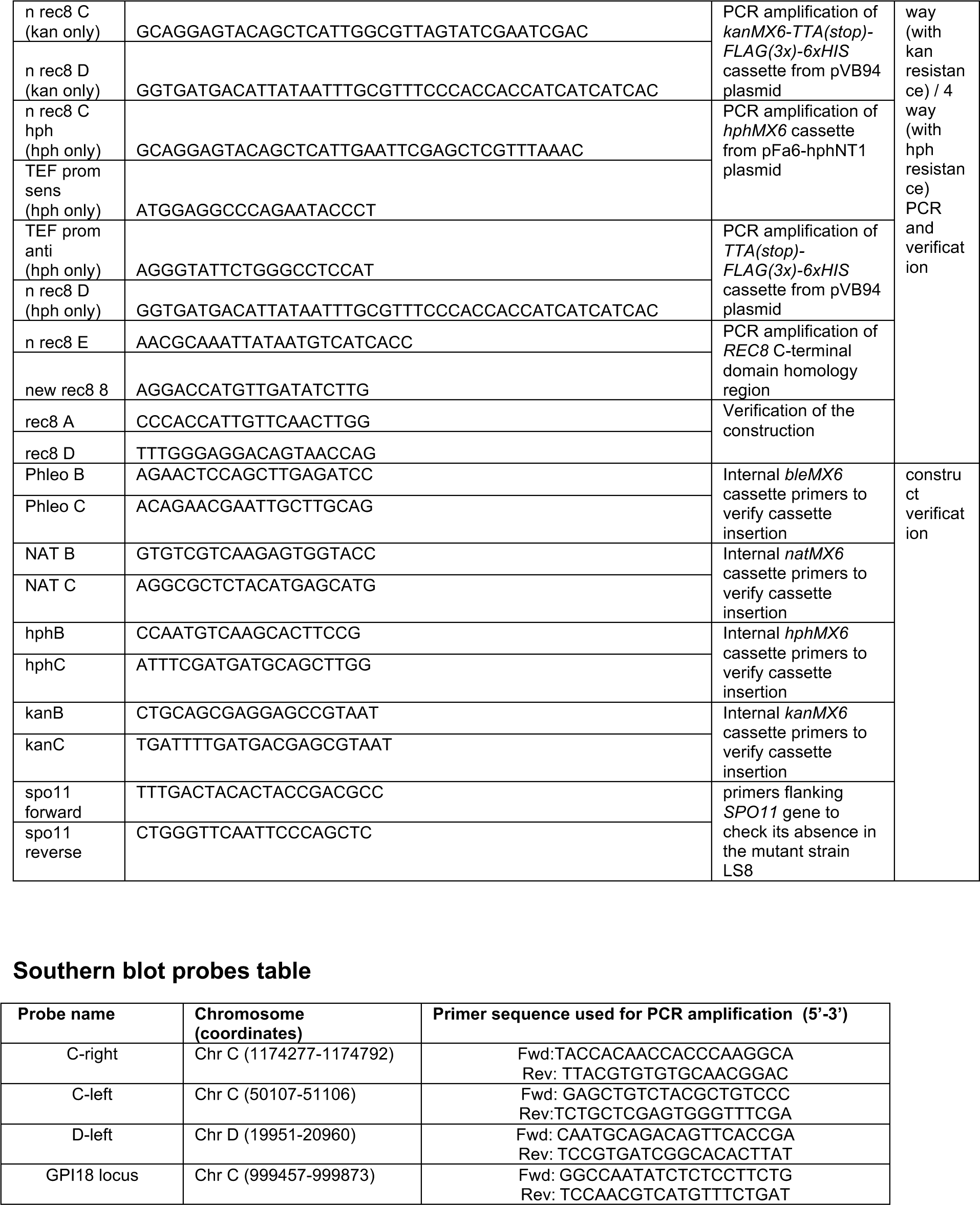
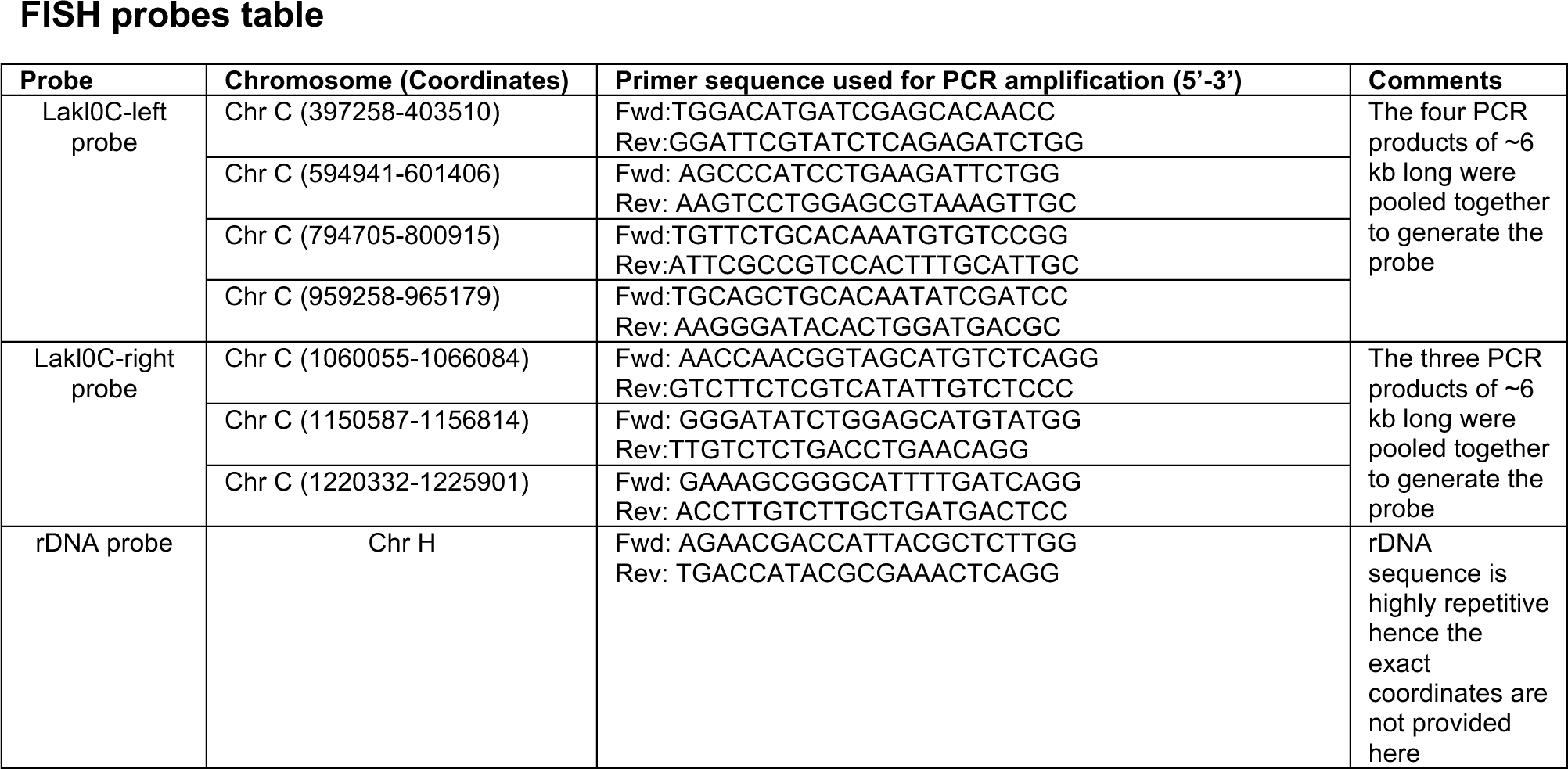
Primers table.

